# Excitatory neurons as the default fate in bifurcation of excitatory and inhibitory neuron lineages

**DOI:** 10.1101/2025.10.29.685327

**Authors:** Mei Wang, Xiao-Ying Qiu, Yun-Bin Duan, Xia Tang, Qi Lu, Yan Li, Hui-Wen Qin, Xing-Tao Long, Meng-Meng Jin, Peng-Yu Fan, Hui Zhang, Ya-Nan Li, Jie He

**Affiliations:** Institute of Neuroscience, Center for Excellence in Brain Science and Intelligence Technology, Chinese Academy of Sciences, Shanghai 200031, China; University of Chinese Academy of Sciences, Beijing, China

## Abstract

In the brain, a precise excitation-inhibition balance underpins every neural operation. Strikingly, across the vertebrate CNS, excitatory and inhibitory neurons often emerge from common progenitors (termed EX-IN lineage), yet how such dichotomous fates diverge has remained enigmatic. Here, exploiting spinal v2a (excitatory)–v2b (inhibitory) and retinal bipolar (excitatory) - amacrine (inhibitory) lineages as EX-IN lineage paradigm, we report a single, conserved genetic program: excitatory neural fate is the default state when sibling-cell communication is silenced; progenitor- and excitatory neuron-expressed homeobox genes actively repress the inhibitory program, while Notch signaling releases this repression to unleash inhibitory identity. Thus, we define a unifying principle governing EX-IN lineage divergence that generate the fundamental dichotomy of neural circuits.

## Main Text

Neurons are functionally categorized as excitatory or inhibitory based on their neurotransmitter profiles. The balance between these two classes is essential for proper neural circuit operation. In the developing vertebrate CNS, excitatory and inhibitory neurons frequently originate from separate developmental lineages. Yet, certain conserved neurogenic programs generate both types from a common progenitor, referred to here as excitatory-inhibitory (EX–IN) lineages. For example, spinal p2 progenitors simultaneously produce glutamatergic v2a (excitatory) and GABAergic v2b (inhibitory) interneurons in zebrafish and mice (*1, 2*). Likewise, a defined lineage in the zebrafish retina gives rise to glycinergic amacrine cells (ACs, inhibitory) and glutamatergic OFF-bipolar cells (BCs, excitatory) as sibling neurons (*3*). Similar bifurcating lineages are found in the mouse cerebellum and hypothalamus (*4, 5*), and even in the human cortex (*6, 7*), highlighting their widespread occurrence and functional importance. However, the developmental mechanism underlying this dichotomy—whether governed by cell-intrinsic factors or extrinsic signals—remains unclear. In particular, is fate bifurcation driven by the asymmetric partitioning of intrinsic determinants during mitosis, or by extrinsic communication between post-mitotic sibling cells? Addressing this question is crucial for understanding how excitatory and inhibitory fate divergence is regulated and how its disruption may contribute to neurodevelopmental disorders.

Here, we leverage the experimental strengths of zebrafish—external development, optical transparency, fast embryogenesis, and advanced genetic tools—to systematically investigate EX–IN lineage bifurcation. Focusing on spinal v2a-v2b and retinal AC-BC lineages, we combine single-cell ablation, precise lineage tracing, and genetic perturbations to uncover the mechanisms enabling a common progenitor to simultaneously produce functionally opposed neurons. Our findings reveal a conserved molecular logic governing fate bifurcation in excitatory-inhibitory lineages and refine the model of neuronal diversification.

## The excitatory neurons serve as the default cell fate in v2a-v2b lineages

To investigate whether excitatory and inhibitory neuronal fate specification in EX-IN lineages is cell-autonomous or requires communication between sibling cells, we employed a strategy of randomly ablating one sibling precursor cell to eliminate intercellular interaction and tracked the fate of the surviving cell (Fig. 1A). We define precursor cell as the cell shortly after the division and have not matured into a specific neuronal type.

**Fig. 1.**
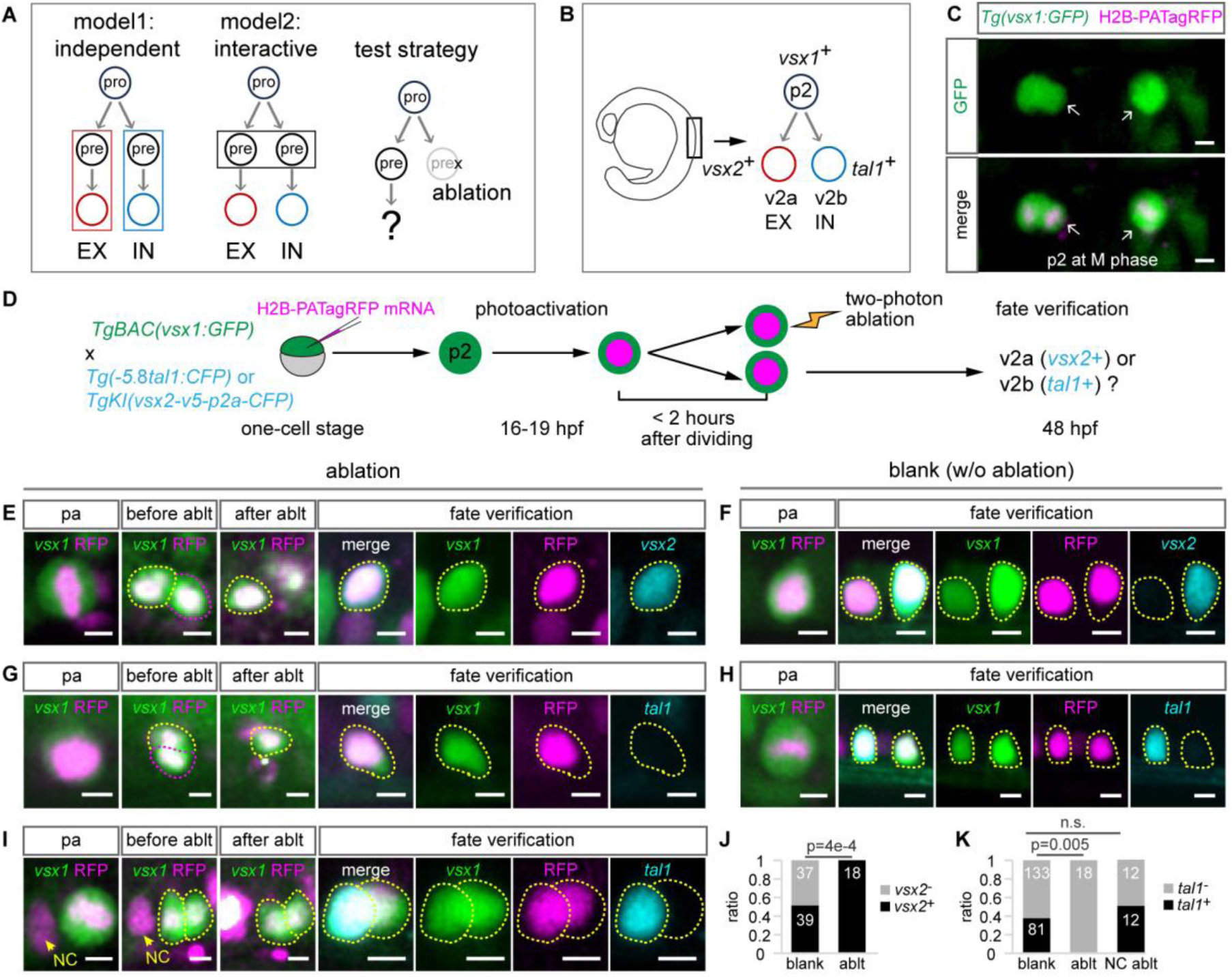
The excitatory neurons are the default cell fate in v2a-v2b lineages. (**A**) Two models of sibling cell fate specification in EX-IN lineages, and the test strategy. (**B**) Schematic of v2a-v2b lineage. (**C**) Label individual p2 cells at M-phase. (**D**) Sibling ablation strategy and fate validation (*vsx2*: v2a marker; *tal1*: v2b marker). (**E** and **F**) Live imaging of v2a fate determination with/without sibling cell ablation. Cell indicated by magenta circle is the ablation target. (**G** and **H**) Live imaging of v2b fate determination with/ without sibling cell ablation. Cell indicated by magenta circle is the ablation target. (I) Live imaging of v2b fate determination with non-sibling neighbor cell (NC) ablation. (J) V2a (*vsx2*^+^) ratio in p2-derived lineages with/without sibling cell ablation. (**K**) V2b (*tal1*^+^) ratio in p2-derived lineages without ablation, with sibling or non-sibling neighbor cell ablation. Numbers showed on boxplots indicate cell numbers. Scale bars: 5 μm. Significance was determined by χ² test.

We used the well-characterized zebrafish v2a-v2b lineage in the spinal cord as the study model (Fig. 1B). The *Tg(vsx1:GFP)* reporter line was used to label p2 progenitors and their derived cell types (*1*). This line exhibits discrete spatial distribution of individual p2 cells, enabling high-fidelity single-lineage tracking (Fig. 1C). To identify v2a and v2b neuronal fate, we engineered and validated reporter lines targeting *vsx2* (*TgKI(vsx2-v5-p2a-CFP*)) and *tal1* (*Tg(−5.8tal1:CFP)*), respectively (fig. S1, A-D). We introduced nuclear localized PATagRFP (photoactivatable TagRFP), a fluorescence protein turns from dark to red with UV stimulation (*8*), into double-transgenic embryos (the crossing of *Tg(vsx1:GFP)* with either *TgKI(vsx2-v5-p2a-CFP)* or *Tg(−5.8tal1:CFP)*) by mRNA injection. To label single p2 progenitors, we photoactivated PATagRFP in individual M-phase p2 cells, identified by their rounded morphology and strong GFP expression (Fig. 1C), at 16-19 hours post fertilization (hpf)—a developmental window, during which many p2 progenitors enter mitosis. Within 2 hours after cytokinesis, we performed random sibling-cell ablation using two-photon microscopy across all spatial coordinates (fig. S2, A–D). Notably, p2 progenitor division axes exhibited variable orientations, as previously described (*1*). Fate specification of surviving cells was assessed at 48 hpf, a time point when *vsx2* and *tal1* reporter signals had reached detectable thresholds (Fig. 1D). In control experiments, we labeled individual p2 cells and tracked the fate of their progeny without ablation, establishing a baseline for unperturbed lineage progression.

In unperturbed control experiments, 37/38 lineages produced one *vsx2*^+^ and one *vsx2*^-^ cell, while only 1 lineage generated two *vsx2*^+^ cells, resulting in a *vsx2*^+^ cell ratio of 0.51 (n = 15 fish; Fig. 1, F and J). In stark contrast, when one sibling cell was ablated within two hours after division, all surviving cells adopted the v2a fate (*vsx2*^+^; 18/18 lineages from 11 fish), a frequency significantly higher than that in controls (p = 4e-4; Fig. 1, E and J). Concomitantly, no surviving cells expressed *tal1*, the v2b marker (0/18 lineages from 15 fish; Fig. 1, G and K). Among 107 control lineages assessed using the *tal1* reporter, 81 sibling pairs exhibited *tal1*^+^/*tal1*^-^ cell combinations, while 26 were *tal1*^-^/*tal1*^-^, yielding a *tal1*^+^ cell ratio of 0.38 (from 55 fish; Fig. 1, G and K). To exclude injury-induced artifacts, we ablated non-sibling neighbor cells and analyzed p2 lineages using the *tal1* reporter. All 12 lineages examined maintained a *tal1*^+^/*tal1*^-^ sibling pair (from 7 fish), confirming that ablation-induced tissue damage did not alter fate specification (Fig. 1, I and K).

Collectively, these data demonstrate that in the absence of sibling-cell, p2 progenitor-derived progeny exclusively adopt an excitatory v2a fate. This identifies excitatory neurons as the default cell fate within v2a-v2b lineages, with inhibitory fate acquisition requiring non-autonomous signals from sibling cells.

## P2/v2a-shared homeobox genes repress inhibitory v2b fate in the default development pathway

Why is the inhibitory v2b neuron fate suppressed in the default developmental pathway? *Tal1*, a basic helix-loop-helix (bHLH) transcription factor, is both necessary and sufficient for specifying inhibitory v2b fate in the developing spinal cord (*9, 10*). Consistent with prior reports, ectopic expression of *tal1* in early v2a-v2b lineages (driven by *vsx1:gal4*) increased the number of GABAergic cells by 1.8 folds (Fig. 2, A and B). Conversely, CRISPR/Cas9-mediated *tal1* knockout in F0 founders significantly reduced GABAergic cell counts in the v2a/v2b layer, confirming its essential role in v2b differentiation (Fig. 2, C and D). Thus, unraveling the upstream regulatory mechanisms of *tal1* is critical for understanding v2b fate inhibition in progenitor cells.

**Fig. 2.**
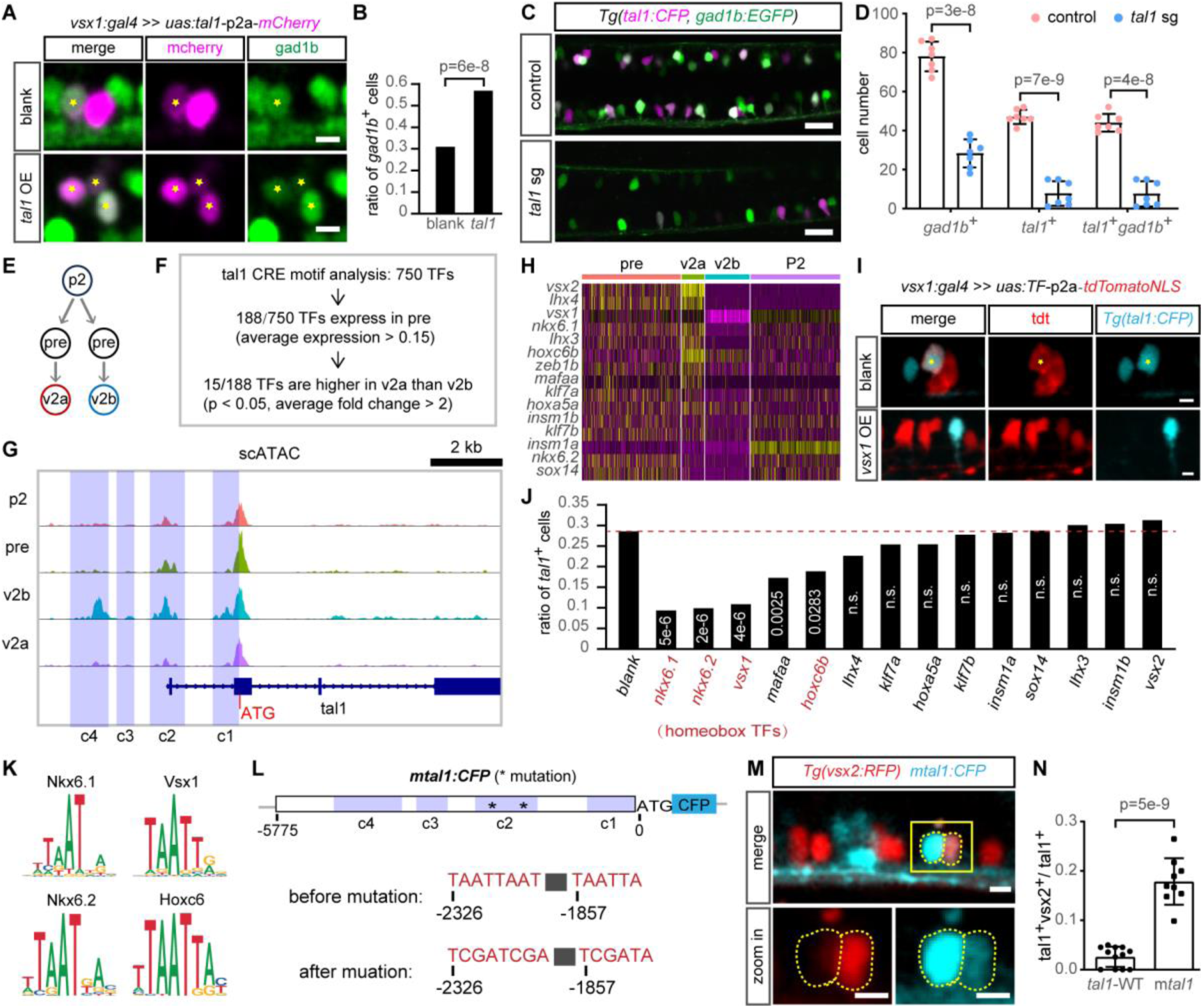
P2/v2a-shared homeobox genes repress inhibitory v2b fate. (**A** and **B**) Effect of *tal1* overexpression (OE) on GABAergic cells: (A) images; (B) Quantification (n = 249 cells in blank, 195 cells in tal1 OE). (**C** and **D**) Effect of *tal1* knockout (by sgRNA injection) on GABAergic cells. (C) images; (D) Quantification, each point stands for one fish. (**E**) V2a-v2b lineage progression schematic. (**F**) Analysis pipeline for identifying *tal1* upstream repressor candidates. (**G**) DNA accessibility in *tal1* coding and promoter regions. Four cis-regulatory elements are defined. (**H**) Expression heatmap of *tal1* repressor candidates from (F). (**I** and **J**) Effect of candidate repressors’ overexpression on *tal1* activation: (I) Images; (J) Quantification (n>170 cells for each group). (**K**) DNA binding motifs of homeobox repressors. (**L**) Mutation of homeobox binding motifs in *tal1* promoter. (**M** and **N**) Activity of mutated *tal1* promoter in v2a: (M) Images; (N) Quantification, each point stands for one fish. Scale bars: 5 μm (A, I, M); 20 μm (C). Significance: χ² test (B, J); t-test (D, N; mean ± SD).

To dissect *tal1*’s upstream regulatory network, we employed single-cell RNA sequencing (scRNA-seq) and single-cell multi-omics (scATAC-seq & snRNA-seq) to characterize molecular signatures during v2a-v2b lineage development. Cells with strong GFP signal were isolated from *Tg(vsx1:GFP)* embryos at 22 hpf for sequencing (fig. S3A). From the total sequenced cells, v2a–v2b lineage cells were subset based on the expression of *vsx1*, *foxn4*, *tal1*, and *vsx2* (fig. S3, B and C and fig. S4, A and B). These cells were segregated into four distinct states according to marker expression: p2 progenitors (*mki67*, *pcna*, *cdk1*), v2a neurons (*vsx2*, *slc17a6a*, *slc17a6b*), v2b neurons (*tal1*, *gata3*, *gad1b*), and intermediate precursors (characterized by low-level expression of progenitor and neuronal markers) (Fig. 2E, fig. S3, D and E, fig. S4, C and D).

Focusing on the 5.8 kb upstream region of *tal1*—a fragment recapitulating endogenous *tal1* activity (fig. S2, B and D)—we identified four cis-regulatory elements (CREs; C1–C4) based on DNA accessibility (Fig. 2G). To uncover *tal1*’s upstream repressors, we predicted transcription factors binding to these CREs based on motif analysis, yielding 750 candidate factors (Fig. 2F). The scRNA-seq analysis revealed 188/750 candidates express in precursor cells (average expression > 0.15), including 15 genes with significantly higher expression in v2a than in v2b cells (Fig. 2, F and H). 14 of these v2a-enriched genes were successfully overexpressed in early v2a-v2b lineages, with *nkx6.1*, *nkx6.2*, *vsx1*, *mafaa*, and *hoxc6b* significantly suppressed *tal1* activation (*tal1*^+^ cell ratios: 9.3%, 9.1%, 10.9%, 17.3%, 18.8% vs. 29% in controls; Fig. 2, I and J). Strikingly, four of the five genes (*nkx6.1*, *nkx6.2*, *vsx1*, *hoxc6b*) are homeobox transcription factors sharing a conserved TAATTA core DNA-binding motif (Fig. 2K). Notably, these homeobox genes start expression in p2 cells (average expression > 0.15), supporting their role in repressing *tal1* at early stage of lineage progression (Fig. 2H).

To test whether relieving repression by these p2/v2a-shared homeobox genes activates *tal1*, we mutated two critical TAATTA motifs in the C2 region of the *tal1* promoter (*11*) (Fig. 2, K and L). Using a mutated *tal1* promoter reporter (*mtal1:CFP*), we observed that 17% of CFP^+^ cells co-expressed the v2a marker *vsx2*, compared to only 3% in wild-type reporters (*−5.8tal1:CFP*), demonstrating *tal1* activation upon homeobox repressors removal (Fig. 2, M and N). Additionally, *tal1* knockout reduced wild-type *tal1* promoter reporter activity, indicating *tal1* autoregulation (Fig. 2, C and D). Therefore, the 17% v2a labeling ratio in mutated reporters was likely an underestimation due to the lack of Tal1 protein in v2a neurons.

To summarize, in the default developmental trajectory of progenitor cells, transcription of the v2b fate determinant *tal1* is directly repressed by multiple p2/v2a-shared homeobox genes, notably *vsx1*, *nkx6.1*, and *nkx6.2*. Disrupting this repressive network enables *tal1* transcription, unveiling a molecular switch for inhibitory v2b fate specification.

## Homeobox repressors are directly suppressed by Notch signaling pathway during inhibitory v2b specification

How do one of the precursor cells adopt v2b fate through sibling cell interactions? It has been well established Notch signaling pathway, a well characterized mediator of lateral inhibition, is asymmetrically activated in v2a-v2b lineages and promotes v2b fate specification (*12–15*). Consistent with this, scRNA-seq analysis revealed that Notch ligands *dll4* and *dlc* are highly expressed in v2a neurons, whereas Notch receptors *notch1a*, *notch1b*, and *notch3* are enriched in v2b cells (foldchange > 1.5; Fig. 3A). Among probable Notch effector genes (Hes/Hey family), *her15.1*, *her6*, and *her4.1* and *her9* show preferential expression in v2b vs. v2a neurons (foldchange > 1.5; Fig. 3A). This asymmetric Notch activity was validated using the *Tg(TP1:h2b-mCherry)* reporter line, supporting a model of lateral inhibition-mediated v2b fate specification (Fig. 3B). *Her15.1*, a Notch effector gene previously linked to v2b induction(*16*), is highly expressed during v2b formation but absent in v2a neurons (Fig. 3, A and C). Ectopic *her15.1* expression in early v2a-v2b lineages significantly expanded v2b neuron proportions (43% vs. 26% in controls; Figures 3D and 3E). Co-expression assays showed that forced *vsx1*, *nkx6.1*, or *nkx6.2* expression abrogated *Her15.1*-mediated *tal1* activation (Fig. 3, F-I), indicating that inhibition of these p2/v2a-shared homeobox repressors is essential for Her15.*1*-driven v2b specification. CUT&Tag analysis further demonstrated Her15.1 chromatin occupancy at regulatory regions of *vsx1*, *nkx6.1*, and *nkx6.2*, providing evidence of direct transcriptional repression (Fig. 3J). This result is consistent with previous study that Her15.1’s repressive function is essential for v2b fate induction (*16*).

**Fig. 3.**
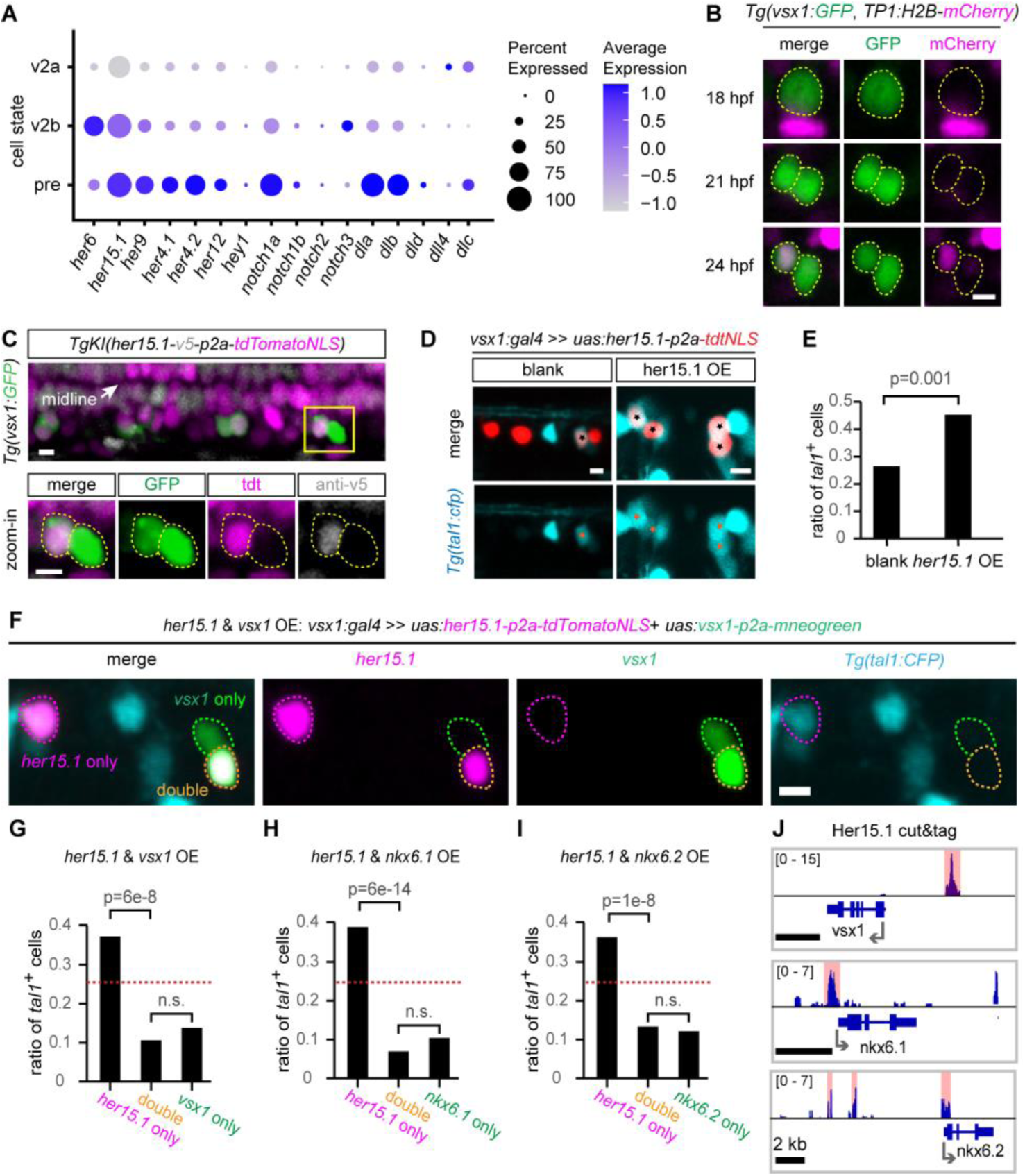
Homeobox repressors are directly suppressed by Notch in inhibitory v2b specification. (**A**) Expression of Notch receptors, ligands and putative effector genes (her/hey proteins) from scRNAseq. (**B**) Time-lapse imaging of Notch activity dynamics (reported by TP1). (**C**) Her15.1 protein distribution in v2a and v2b neurons. Below panels show the zoom-in image in the above yellow rectangle. (**D** and **E**) Effect of *her15.1* overexpression on *tal1* activation: (D) images; (E) Quantification (n = 167, 117 cells for blank and her15.1 OE, respectively). (**F**) Images of her*15.1* & *vsx1* co-overexpression. (**G**-**I**) Quantification the effects of *her15.1* & *vsx1* (G), *her15.1* & *nkx6.1* (H), *her15.1* & *nkx6.2* (I) co-overexpression on *tal1* activation (n > 124 cells for each group/boxplot). Red dot lines indicate the blank value refers to (E). (**J**) Her15.1 protein binding frequency around DNA locus of homeobox repressors. Binding peaks are indicated. Scale bars: 5 μm. Significance: χ² test.

Collectively, Notch signaling is robustly activated in v2b but silenced in v2a lineages. The Notch effector Her15.1 directly inhibits p2/v2a-shared homeobox repressors (*vsx1*, *nkx6.1*, *nkx6.2*), derepressing *tal1* transcription. In sibling precursor cells with elevated Notch activity, this regulatory cascade activates *tal1*, redirecting differentiation from the default excitatory v2a toward inhibitory v2b fate.

## Homeobox genes shared by progenitors and excitatory bipolar cells repress inhibitory sibling neuronal fate

To explore evolutionary conservation of EX-IN lineage regulatory mechanisms, we investigated neural differentiation in zebrafish retinal AC-BC lineages. AC-BC lineages derive from *atoh7⁺vsx1⁺* progenitors during terminal division (*3*). Leveraging FACS-sorted *atoh7*⁺ cells for single-cell multiomics (fig. S5, A-G) and reanalyzing published scRNA-seq data (*3*) (fig. S6, A-G), we identified four distinct cell states in AC-BC lineage progression: (progenitor, precursor, AC, BC) according to the established marker gene expression (Fig. 4A and fig. S5, F-G and fig. S6, F-G).

**Fig. 4.**
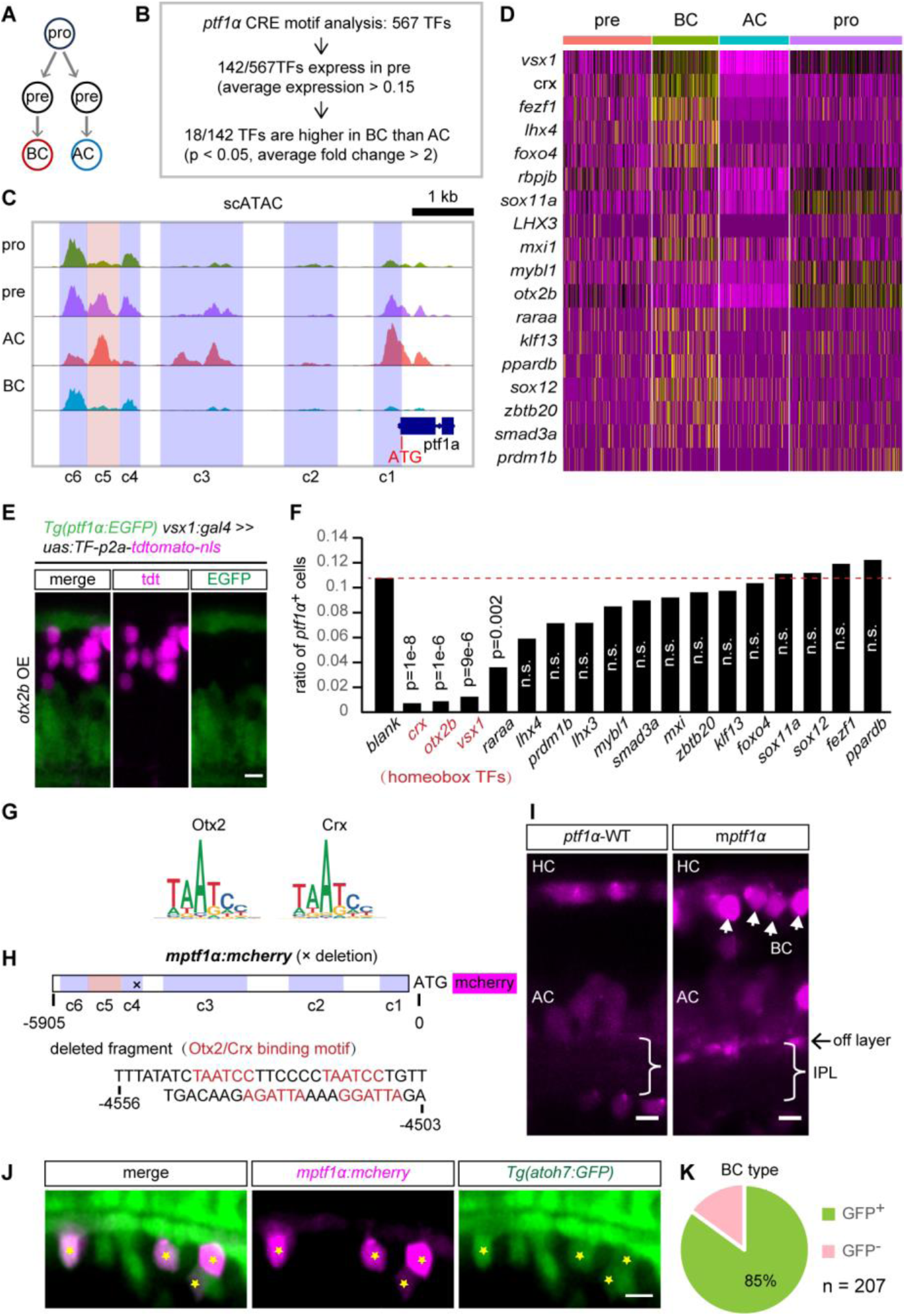
Progenitor/BC-shared homeobox genes repress inhibitory AC fate. (**A**) AC-BC lineage progression schematic. (**B**) Analysis pipeline for identifying *ptf1α* (AC fate determinant) upstream repressor candidates. (**C**) DNA accessibility in *ptf1α* coding and promoter regions. Six cis-regulatory elements are defined. (**D**) Expression heatmap of *ptf1α* upstream repressor candidates from (C). (**E** and **F**) Effect of candidate repressors’ overexpression on *ptf1α* activation: (E) Images; (F) Quantification (n>300 cells for each group). (**G**) DNA binding motifs of Otx2 and Crx. (**H**) Deletion of Otx2/Crx binding motifs in *ptf1α* promoter. (**I**) The reporter signal of wild-type and mutated *ptf1α* promoter. (**J** and **K**) The BC subtype labeled by mutated *ptf1α* promoter (*atoh7* labels off-subtype BC): (J) Representative images; (K) Quantification. Scale bars, 5 μm. Significance: χ² test

The bHLH transcription factor *ptf1α* acts as the master regulator of AC fate specification, being both essential and sufficient for AC differentiation(*17, 18*). A 5.9 kb promoter region upstream of the *ptf1α* start codon recapitulates its endogenous expression pattern, as validated by the strong concordance between its reporter activity and bacterial artificial chromosome (BAC)-driven reporter signals (Fig. S5H). In silico analysis identified six cis-regulatory elements (C1–C6) within this 5.9 kb promoter, guided by chromatin accessibility profiles across AC–BC lineage development (Fig. 4C). Motif analysis predicted binding sites for 567 candidate transcription factors at these elements. Among these, 142 were positively expressed in precursor cells (average expression > 0.15), including 18 factors significantly enriched in BCs versus ACs (p < 0.05, average fold-change > 2; Fig. 4, B and D). We ectopically expressed 17/18 BC-enriched transcription factors in early AC-BC lineages (driven by *vsx1:gal4*). Overexpression of *crx*, *otx2b*, *vsx1*, and *raraa* elicited significant repression of *ptf1α* activity (*ptf1α*^+^ cell percentage: 0.7%, 0.9%, 1.2%, 3.6% versus 10.8% in blank controls; Fig. 4, E and F). Notably, three of the four repressors (*crx*, *otx2b*, and *vsx1*) are homeobox genes that most potently suppress *ptf1α* transcription. These homeobox genes exhibit high expression in progenitor cells (Fig. 4D), supporting their role in repressing AC fate commitment during early stages of lineage progression.

We next investigated whether the removal of this repression could activate *ptf1α* expression in BCs. To address this, we deleted a 54-bp fragment containing four Otx2b/Crx binding motifs (GGATTA/TAATCC) within the C4 regulatory region—a genomic segment that is chromatin-accessible in progenitors, precursor cells and BCs (Fig. 4, C and G and H). As anticipated, the mutated *ptf1α* promoter ectopically directed reporter signal (*mptf1α:mCherry*) in BCs (Fig. 4I). BCs are categorized into off-, on-, and on-off subtypes based on the stratification of their axons within the inner plexiform layer (IPL)(*19*). Notably, the reporter signal driven by the mutated *ptf1α* promoter was specifically enhanced in the IPL sublamina corresponding to the off layer (Fig. 4I). Further molecular characterization revealed that 85% of these labeled BCs are *aoth7*^+^ (n=207; Fig. 4, J and K), a marker that discriminates off-BCs from non-off subtypes(*3*).

Together, progenitor/BC-shared homeobox genes *crx*, *otx2b*, and *vsx1* repress the AC fate determinant *ptf1α*. Disrupting Otx2b/Crx binding in the *ptf1α* promoter suffices to activate *ptf1α* transcription in off-BCs, revealing a conserved mechanism where excitatory fates are maintained by active repression of inhibitory programs.

## Notch directly inhibits homeobox suppressors to drive inhibitory amacrine cell fate

In AC-BC lineages, Notch signaling promotes the differentiation of inhibitory ACs(*20*). Consistent with this role, our scRNA-seq analysis revealed that the Notch ligand *dll4* is preferentially expressed in BCs (foldchange > 1.5), whereas Notch receptors (*notch1a, notch1b, notch3*) are enriched in ACs (Fig. 5A). Multiple putative Notch effector genes from the hes/hey family (*hey1, her15.1, her9, her4.1, her4.2, her12*) showed high expression in ACs (Fig. 5A). Ectopic expression of *hey1*, *her15.1*, or *her12* in early AC-BC lineages potently activated *ptf1α*, increasing the proportion of *ptf1α*^+^ cells from 9.9% to ∼32% (Fig. 5, B and C). To dissect the conserved Notch mechanism, we focused on *hey1* (over *her15.1*) to ensure findings were not isoform-specific.

**Fig. 5.**
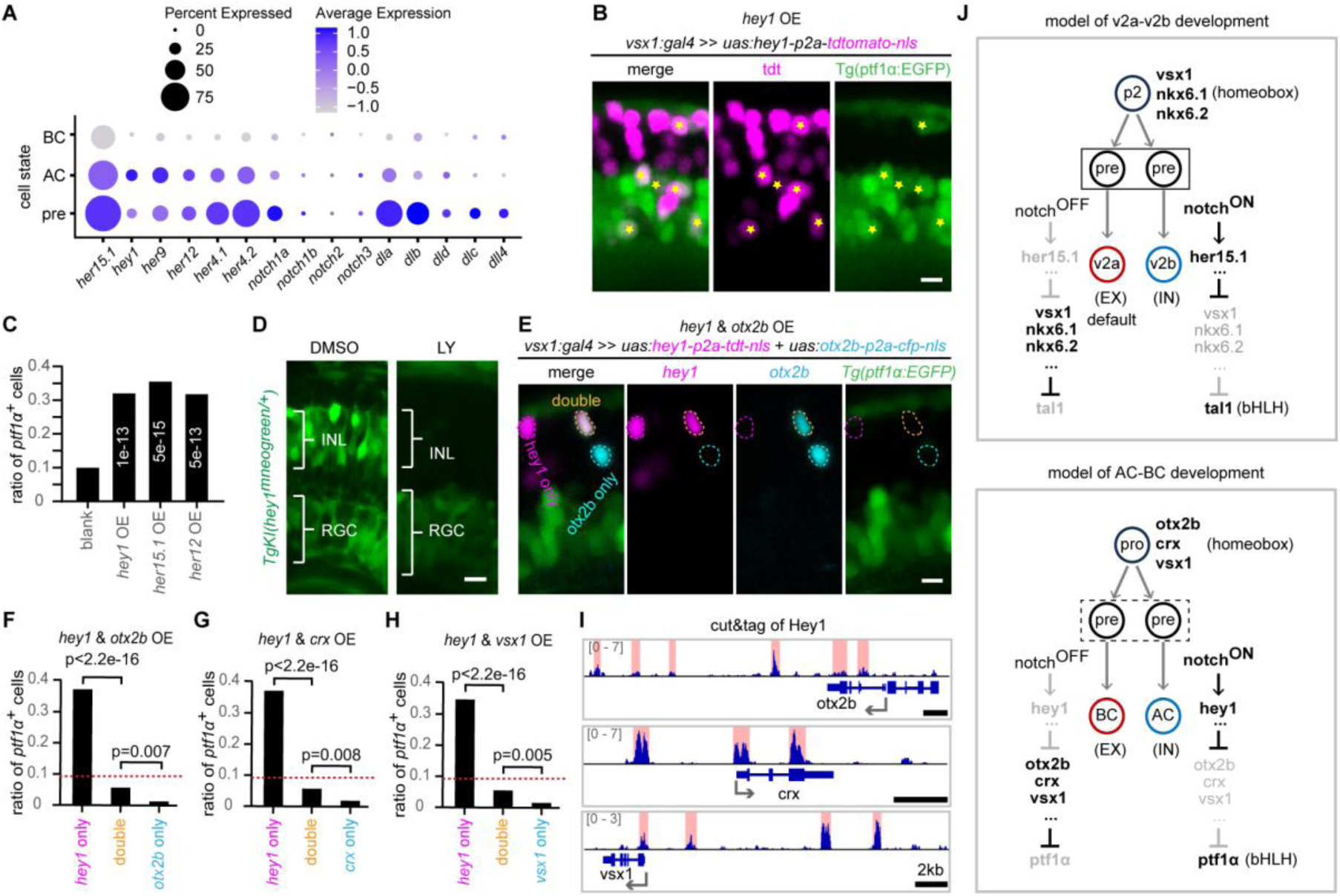
Homeobox repressors are directly suppressed by Notch in inhibitory AC specification. (**A**) Expression of Notch receptors, ligands and putative effector genes (her/hey proteins) from scRNAseq. (**B** and **C**) Effects of *hey1*, *her15.1* and *her12* overexpression on *ptf1α* activation: (B) Representative images; (C) Quantification  n >300 cells for each group). (**D**) The influence of Notch blocker on *hey1* activity in embryonic retina. (**E**) Images of *hey1* & *otx2b* co-overexpression. (**F**-**H**) Quantification the effects of: (F) *hey1* & *otx2b*; (G) *hey1* & *crx*; (H) *hey 1*& *vsx1* co-overexpression (n > 280 cells for each group/boxplot). Red dot lines represent the blank value refers to (C). (**I**) Hey1 protein binding frequency around DNA locus of homeobox repressors. Binding peaks are indicated. (**J**) Summary model. Scale bars: 5 μm. Significance: χ² test.

To validate *hey1* as a Notch effector, we generated a *hey1* reporter line Tg*KI(hey1^mNeonGreen/+^)* in which one allele of endogenous *hey1* is replaced by mNeogGreen fluorescence protein. Treatment with the Notch inhibitor LY411575 abrogated *hey1* reporter signal in the inner nuclear layer (INL), where ACs and BCs reside, confirming *hey1* as a canonical Notch effector in AC-BC development (Fig. 5D).

We next investigated whether *hey1* activates *ptf1α* by relieving transcriptional repression. Co-overexpression of *hey1* with AC-repressing homeobox genes (*otx2b, crx, vsx1*; Fig. 4F) in early AC-BC lineages revealed that while *hey1* alone induced *ptf1α*^+^ cells at 37.2%, co-expression with *otx2b* reduced this to 5.7% (Fig. 5, E and F)—below the blank control (9.9%, the blank of Fig. 5C). Similar suppression occurred with *hey1* & *crx* and *hey1* & *vsx1* co-overexpression (Fig. 5, G and H), demonstrating that *hey1*-mediated *ptf1α* activation requires repression of *otx2b*, *crx*, and *vsx1*. Hey1 Cut&Tag profiling further revealed direct binding of Hey1 to putative cis-regulatory elements of these homeobox repressors (Fig. 5I), establishing direct transcriptional repression.

In summary, Notch signaling is asymmetrically activated in sibling precursor cells of AC-BC lineages. In cells with high Notch activity, the effector Hey1 directly represses AC-repressing homeobox genes (*otx2b*, *crx*, *vsx1*), relieving the AC fate determinant *ptf1α* to drive inhibitory neuronal commitment. Conserved mechanisms likely operate via other Notch effectors (e.g., Her15.1) in these lineages.

## Discussion

The emergence of functionally opposed neurons from shared progenitors across neural tissues implies conserved developmental logic. In the zebrafish v2a-v2b lineage, excitatory neurons constitute the default precursor fate, while inhibitory v2b specification requires sibling-cell interactions. Remarkably, in both the v2a-v2b and AC-BC lineages, a key inhibitory fate determinant from the bHLH family (*tal1* in v2a-v2b; *ptf1α* in AC-BC) is redundantly suppressed by multiple homeobox genes. Notch signaling is asymmetrically activated in sibling cells, likely initiated through lateral inhibition. Meanwhile, Notch signaling, asymmetrically activated in sibling cells, directly relieves homeobox-mediated repression of the inhibitory fate determinant, thereby driving inhibitory neuron production (Fig. 5J).

### Excitatory neurons as the default fate in EX-IN bifurcation

The ablation experiment in v2a-v2b lineages demonstrates that excitatory v2a neurons represent the default fate of precursor cells. The inhibitory fates are redundantly repressed by specific sets of homeobox genes which are expressed from progenitors through to excitatory neurons: *nkx6.1*/*nkx6.2*/*vsx1* in v2a-v2b lineages and *otx2b*/*crx*/*vsx1* in AC-BC lineages. Beyond repressing the inhibitory fate, these homeobox genes may also act as selector genes directly promoting excitatory fates. For instance, *otx2b* promotes BC generation in retina (*3*). Although homeobox genes are central, other genes likely also contribute to promoting excitatory and repressing inhibitory fates. We therefore propose a model wherein a genetic module expressed in progenitor/precursor cells simultaneously promotes excitatory identity and represses the inhibitory alternative, thereby establishing the excitatory fate as the default in EX-IN lineage bifurcations. This model may have broader applicability, as *tal1* and *ptf1α* extensively drive GABAergic neuron generation in hindbrain and spinal cord (*23*). In these regions, homeobox genes (such as patterning genes) likely repress *tal1* or *ptf1α* in precursors, requiring disinhibition for GABAergic specification over glutamatergic fate. This hypothesis is supported by studies in the mouse dorsal spinal cord (*24, 25*). We therefore speculate that excitatory neurons represent the default fate in the bifurcation of excitatory and inhibitory neural fate in this situation. However, the situation may differ in regions where inhibitory neuronal fate is determined by genetic drivers rather than *tal1* or *ptf1α*.

### Sibling-cell interaction establishes fate asymmetry in v2a-v2b lineages

Our findings demonstrate that sibling-cell interaction is essential for driving fate asymmetry in the v2a-v2b lineage. However, we cannot rule out the asymmetric inheritance of fate determinants in the terminal division, which has been widely implicated in asymmetric fate specification(*21, 22*). We showed that—at least in the v2a-v2b system— this intrinsic mechanism alone is insufficient to robustly establish inhibitory neuronal fate. This discovery raises two plausible mechanistic models for how sibling-cell interaction governs lineage segregation: 1. *Asymmetric Inheritance Model*: The P2 division produces sibling precursors with subtle differences due to uneven inheritance of fate determinants. Sibling-cell interaction (e.g. Notch lateral inhibition) then amplifies this initial imbalance, leading to sharply distinct transcriptional and functional outcomes. 2. *Stochastic Symmetry-Breaking Model*: The two sibling precursors are initially equipotent, with stochastic fluctuations in gene expression (e.g. Notch ligands) providing the variation. Sibling-cell interaction subsequently reinforces these fluctuations, driving the cells toward divergent fates. Distinguishing between these models will not only deepen our understanding of EX-IN lineage bifurcation but also reveal broader principles of cell-fate decision-making in development.

## Acknowledgments

We thank Prof. Zilong Wen for Tal1 antibody, Dr. Jiarui Mi for *Tg(TP1:h2b-mcherry)* strain and Dr. Wenjuan Zhang from Weijun Pan’s lab for whole-mount immunostaining assistance. We are grateful to Yuanzhen Zhu for scientific discussions. We acknowledge Qian Hu, Dan Xiang, Xuxing (Optical Imaging Core Facility) for two-photon ablation support and Lijuan Quan, Dr. Min Zhang and Zhenning Zhou (Molecular and Cellular Biology Core Facility) for NGS library preparation.

## Funding

Creative Research Groups of the National Natural Science Foundation of China: 32321003 (JH)

STI2030-Major Projects: 2021ZD0204500 (JH)

National Natural Science Foundation of China: 32471029 (JH)

National Natural Science Foundation of China: 31871035 (JH)

Shanghai Natural Science Foundation: 25ZR1402526 (MW)

National Natural Science Foundation of China: 32500853 (MW)

China Postdoctoral Science Foundation: 2021M703308 (MW)

Shanghai Post-doctoral Excellence Program: 2021400 (WM)

## Author contributions

Conceptualization: MW, JH

Methodology: MW, XYQ, MMJ

Investigation: MW, XYQ, YBD, YL, XTL, QL, XT, YNL

Data curation: MW, HZ

Funding acquisition: MW, JH

Project administration: MW

Supervision: JH

Writing – original draft: MW, JH

## Data and materials availability

The authors declare that all data needed to evaluate the conclusions in this study are available in the main text or the supplementary materials. Raw data of single-cell RNA sequencing, single-cell multiome sequencing, CUT&Tag sequencing reported in this paper have been submitted to the Genome Sequence Archive of National Genomics Data Center, Beijing Institute of Genomics (BIG), Chinese Academy of Sciences (http://gsa.big.ac.cn/) with accession number CRA028101. The code, detailed analysis records and processed data are available at Zonodo: https://doi.org/10.5281/zenodo.16261575. Plasmids and zebrafish lines made in this study will be distributed upon request to other investigators under a Material Transfer Agreement.

## Materials and Methods

### Zebrafish husbandry

Zebrafish lines were bred and maintained at 28 ℃ on 14-hour-light/10-hour-dark cycles. Embryos were obtained from natural spawning and raised at 28 ℃ in embryo medium (NaCl 5.03 mM, KCl 0.17 mM, CaCl_2_•2H_2_O 0.33 mM, MgSO_4_•7H_2_O 0.33 mM, Methylene blue 0.0002% (w/v)) unless otherwise indicated. Embryos were staged by hours post-fertilization (hpf) before 72 hpf as previously described (*26*) and days post-fertilization (dpf) after that. To prevent pigmentation, embryos designated for imaging were treated with 0.003% phenylthiourea (PTU, Sigma-Adrich, Cat# P7629) beginning at 8 hpf. As zebrafish sex cannot be determined before 25 dpf, the sex of the experimental animals was unknown. All animal procedures performed in this study were approved by the Animal Use Committee of Center for Excellence in Brain Science and Intelligence Technology, Chinese Academy of Sciences (NA-069-2023).

### Zebrafish transgenic lines

The following published transgenic lines were used: *Tg(vsx1:GFP)*(*1*), *Tg (vsx2:DsRed)*(*27*), *Tg(ptf1α:EGFP)*(*28*), *Tg(atoh7:GFP)*(*29*), *Tg(gad1b:EGFP)*(*3*), *Tg(TP1:h2b-mcherry)*(*30*), *Tg(atoh7:gapRFP)*(*31*), *Tg(atoh7:turboGFP-dest1)*(*3*), *Tg(uas:kaede)*(*32*). The following transgenic lines were generated in this study: *Tg(−5.8tal1:cfp)*, *Tg(−5.9ptf1α:mCherry)*, *Tg(mptf1α:mCherry)*, *Tg(uas:her15.1-v5-p2a-tdTomatoNLS)*, *Tg(vsx1:gal4)*, *Tg(lhx3:gal4)*, *TgKI(vsx2-v5-p2a-CFP)*, *TgKI(her15.1-v5-p2a-tdTomatoNLS)*, *TgKI(hey1^mneogreen^)*, *TgKI(hey1-v5-p2a-gal4)*.

### Plasmid construction

To construct *−5.8tal1:CFP* and *-5.9ptf1α:mCherry*, a 5.8 kb fragment upstream of *tal1* and a 5.9 kb upstream of ptf1α start codon were cloned from zebrafish genomic DNA, respectively. *Mtal1: CFP* and *mptf1α:mCherry* were generated from *−5.8tal1:CFP* and *- 5.9ptf1α:mCherry*, respectively, by introducing specific mutations via site-directed mutagenesis with the indicated primers (table S1). For *pCS2:H2B-PATagRFP* construction, the PATagRFP DNA sequence was obtained from Addgene (https://www.addgene.org/) and synthesized (GENEWIZ). The H2B coding region was cloned 48 hpf zebrafish cDNA. Both H2B and PATagRFP fragments were then inserted into the pCS2 vector. *Uas:her15.1-v5-p2a-tdTomatoNLS* was generated from the *UAS:otx2-p2a-tdTomatoNLS* (*3*), by replacing the *otx2* coding sequence with a *her15.1-v5* cassette. This cassette was constructed by fusing a 3xV5 tag to the C-terminus of Her15.1. (NLS denotes nuclear localization signal). All uas:TF-p2-FP plasmids (e.g. *uas:tal1-p2a-tdTomatoNLS*) were derived from *uas:otx2-p2a-tdTomatoNLS* (*3*), by replacing either the *otx2* coding region or the *tdTomatoNLS* sequence with the target transcription factor or fluorescent protein coding sequence, respectively. DNA fragments flanked by ∼17 bp homologous arms were assembled using the ClonExpress MultiS One Step Cloning Kit (Vazyme, Cat# TD-903).

The BAC plasmid *lhx3:gal4* were generated as previously described (*33*). Briefly, CH73-365E20 bacteria harboring the lhx3 BAC were obtained commercially (Source BioScience). BAC DNA was extracted using the NucleoBond BAC 100 kit (MACHEREY-NAGEL, Cat# 740579) according to the manufacturer’s instructions and electroporated into SW105 bacteria. A cassette containing ∼50-bp homology arms, *gal4*, a polyadenylation signal, and a kanamycin resistance gene (*neoR*) was inserted into the *lhx3* start codon via recombineering. The FRT-*neo*-FRT cassette was subsequently excised by L-arabinose induction. Primers used for plasmid construction are listed in table S1.

### Transgenic fishline generation

The transgenic strains *Tg(−5.8tal1:CFP)*, *Tg(−5.9ptf1α:mCherry)*, *Tg(mptf1α:mCherry)*, *Tg(uas:her15.1-p2a-tdTomatoNLS)*, *Tg(lhx3:gal4)* and *Tg(vsx1:gal4)*(*3*) were created with Tol2 system (*34*), by co-injecting 10 ng/μl plasmids and 50 ng/μl tol2 mRNA into wild-type embryos at 1-cell stage. Injected embryos were screened for positive founder.

The transgenic knock-in lines *TgKI(vsx2-v5-p2a-CFP)*, *TgKI(her15.1-v5-p2a-tdTomato-NLS)*, *TgKI(hey1^mneogreen^)*, *TgKI(hey1-v5-p2a-gal4)* were generated using Tild-CRISPR technology (*35*). Briefly, donor plasmids containing 200-1000 bp homology arms were constructed. To prevent Cas9 cleavage, synonymous mutations were introduced into the sgRNA target sites within the donor plasmids. Donor fragments, comprising the homology arms and the insertion cassette, were PCR-amplified from these plasmids. Embryos at the early one-cell stage were co-injected with donor fragments (20 ng/μl), Cas9 protein (400 ng/μl, Novoprotein, Cat# E365-01A), and sgRNA (200 ng/μl). Injected embryos were screened for positive founders, and established lines were validated by sequencing. In the *TgKI(vsx2-v5-p2a-CFP)*, *TgKI(her15.1-v5-p2a-tdTomatoNLS)*, *TgKI(hey1-v5-p2a-gal4)* lines, the insertion cassettes (v5-p2a-CFP, v5-p2a-tdTomatoNLS, v5-p2a-gal4, respectively) were in-frame fused to the C-terminal of target genes. In *TgKI(hey1^mneogreen^)*, one *hey1* allele was replaced by the mNeoGreen coding sequence. Primers used for knockin fishline construction are listed in fig. S5.

### sgRNA and mRNA synthesis

sgRNAs were designed using the CRISPRScan online tool (*36*). Templates for sgRNA *in vitro* transcription were generated by PCR-amplifying the standard sgRNA scaffold fragment (*37*) using specific forward primers and a universal reverse primer (table S1). sgRNAs were synthesized *in vitro* using the MEGAshortscript T7 kit (Invitrogen, Cat# AM1354) according to the manufacturer’s instructions. To synthesize H2B-PATagRFP mRNA, the *pCS2:H2B-PATagRFP* plasmid was linearized with NotI-HF (NEB, Cat# R0189V) and used as a template for *in vitro* transcription with the mMESSAGE mMACHINE SP6 Kit (Invitrogen, Cat# AM1340) following the manufacturer’s protocol.

### Gene knockout and overexpression

To knock out tal1 in G0, an equimolar mixture of four sgRNAs targeting the *tal1* coding region (total 200 ng/μl) and Cas9 protein (400 ng/μl, Novoprotein, Cat# E365-01A) was coinjected into *Tg(−5.8tal1:CFP, gad1b:EGFP)* embryos at one-cell stage (*38*). Control embryos were injected with scramble sgRNA (*39*). Embryos were imaged at 48 hpf.

To overexpress transcription factors with gal4/UAS system (*40*) in v2a-v2b or AC-BC lineages, we injected plasmids *uas:TF-p2a-FP* (15 ng/μl), BAC plasmids (*3*) (15 ng/μl) and Tol2 transposase mRNA (50 ng/μl) into hybrid embryos derived from *Tg(vsx1:gal4)* and fate reporter line (e.g., *Tg(−5.8tal1:CFP)*). Coinjection of the *vsx1:gal4* BAC plasmid enhanced overexpression efficiency. Positive embryos were imaged on confocal at 48 hpf (v2a-v2b lineages) or at 4 dpf (AC-BC lineages).

### Whole-mount immunostaining

To evaluate the efficacy of *Tg(−5.8tal1:CFP)*, *Tg(−5.8tal1:CFP)* embryos at 24 hpf were de-chorionized and fixed by 4% paraformaldehyde overnight at 4 ℃. Dehydrate sample stepwise in 25%, 50%, 75%, 100%, 100% methanol/PBST ((PBS containing 0.1% Tween-20; 10 minutes for each step) and store overnight in methanol at -20 ℃. Rehydrate the sample stepwise in 75%, 50%, 25% methanol/PBST (10 minutes for each step). Wash sample gently three time with PBST. Treat sample with acetone (pre-chilled at -20 ℃) for 30 minutes at -20 ℃. Wash sample gently three time with PBST, 5 minutes for each. Block the sample in Block Solution (PBS with 0.3% TritonX-100, 1% DMSO, 1% BSA, 10% donkey serum) for 1 hour at RT. Dilute the first antibodies with Incubating Buffer (PBS with 0.3% TritonX-100, 1% DMSO, 1% BSA, 2% donkey serum), incubate sample with diluted first antibody overnight at 4 ℃. The primary antibodies were: Rabbit anti-Tal1 (1:50) (*41*), Chicken monoclonal anti-GFP (1:2000, Proteintech Group, Cat# 50430–2-AP). Wash sample four time with Incubating Buffer, 30 minutes for each. Dilute the second antibodies with Incubating Buffer and incubate the sample with diluted second antibody overnight at 4 ℃. The secondary antibodies were: AlexaFluor 647 Donkey anti-Rabbit (1:1000), AlexaFluor 488 Donkey anti-Chicken (1:1000). Wash sample four time with Incubating Buffer, 30 minutes for each.

### Confocal imaging

Embryos at desired stages were anesthetized by 0.04% Tricaine methanesulfonate (MS-222, Sigma-Adrich, Cat# E10521), embedded in 1% low-melting agarose (Sigma-Adrich, Cat# A0701), imaged using the inverted laser-scanning confocal microscope (FV1200, Olympus). V2a-v2b lineages in spinal cord were imaged from dorsal view under 30× (oil, NA= 1.05) objectives. AC-BC lineages in retina were imaged from lateral view under 60× (water, NA=1.20) objectives.

### Cell ablation

Hybrid embryos from *Tg(vsx1:GFP)* and *TgKI(vsx2-v5-p2a-CFP)* (or *Tg(−5.8tal1:CFP)*) crosses were injected with H2B-PATagRFP mRNA (250 ng/μl) at the one-cell stage. To delay development for convenient imaging, injected embryos were maintained at 26.5 °C. Embryos exhibiting GFP signal were selected at the ∼18-somite stage based on morphology (*26*). Selected embryos were dechorionated, anesthetized in 0.04% MS-222 (Sigma-Adrich, Cat# E10521), and embedded belly-up in 0.6% low-melting agarose (six embryos for each batch). Samples were incubated in embryo medium containing 0.02% MS-222 and 0.003% PTU (Sigma-Adrich, Cat# P7629). P2 cells in metaphase (identified by strong GFP signal and rounded morphology) were photoactivated using UV light on an inverted confocal microscope (Olympus FV1200, 30x oil objective, NA=1.05). P2 labeling for each batch required <1 hour. Following photoactivation, agarose blocks containing embryos were carefully inverted (for upright two-photon microscopy), excess liquid was removed, and blocks were immobilized with a drop of melted agarose. Incubate the samples again into embryo medium (with 0.02% MS-222, 0.003% PTU). Imaging was performed on an upright two-photon microscope (Bruker) with a 20x water objective (Olympus, NA=1.0), simultaneously detecting GFP and RFP by laser (Coherent Chameleon Discovery) at 1040 nm excitation. The focal plane was adjusted to visualize the target cell while maximize the distance to the sibling cell, minimizing non-specific ablation. Target cells were imaged pre- and post-ablation at 3x zoom. Ablation was performed at 1040 nm using a single 200 ms laser pulse. Laser power (400-700, sample-dependent) was calibrated in a non-target region near the target cell prior to ablation. Ablation was repeated as necessary until the target region showed morphological defects (appearance of bright dots) or the absence of the target cell’s GFP signal. After ablation, embryos were individually released into a 24-well plate containing embryo medium with 0.003% PTU and maintained at 28 °C. Surviving sibling cell fate was assessed via confocal imaging at 48-54 hpf.

### Preparation of single-cell suspension

single-cell suspension was prepared as previously described (*39*). Specifically, samples were dissociated in 200 μl papain solution at 37°C for 15 minutes. Papain solution contains 4 μl of papain (Worthington Biochemical Corporation, Cat# LS003126), 4 μl of 1% DNase I (Sigma-Adrich, Cat# DN25), 8 μl of 12 mg/ml L-cysteine in 184 μl DMEM/F12 (Gibco, Cat# 11330032). The dissociation reaction was terminated by adding 800 μl of washing buffer. The dissociation was terminated and pipetted using 800 μl washing buffer (1 ml washing buffer was prepared by adding 6.5 μl 45% glucose, 5 μl 1M HEPES, 50 μl fetal bovine serum, 938.5 μl DPBS). The resulting suspension was filtered through a 40 μm cell strainer and centrifuged at 500 *g* for 5 min at 4°C. The supernatant was discarded, and the cell pellets was resuspended in PBS containing 0.04% bovine serum albumin (BSA).

### Single-cell RNA sequencing library construction and data analysis

For v2a-v2b lineages, embryonic trunks (heads and yolk sacs removed) from *Tg(vsx1:GFP)* embryos at ∼20 hpf (determined by morphology)(*26*) were collected. Single-cell suspension was prepared as described above. Cells with strong GFP signal were isolated by fluorescence-activated cell sorting (FACS; BD FACSAria Fusion; fig. S3A). The sorted cell suspension was centrifuged at 500 g for 5 minutes at 4 °C, and the pellet was resuspended in 30 μl PBS-0.04% BSA. Single-cell libraries were then generated using the Chromium Single Cell 3’ Library & Gel Bead Kit (10x Genomics, Cat# PN-120237) following the manufacturer’s protocol. Libraries were sequenced on an Illumina NovaSeq 6000 platform with 150 bp paired-end reads.

Raw sequencing data were processed using Cell Ranger software (v7.1.0) to perform sample demultiplexing, align reads to the zebrafish reference genome (GRCz11), and generate a gene expression count matrix. Subsequent analysis, including quality control, normalization, dimensionality reduction, clustering, and differential expression analysis, was performed using the Seurat R package (v5.1.0). For AC-BC lineages, we reanalyzed a published single-cell RNA sequencing dataset of *atoh7*^+^ PRCs (*3*).

### Single-cell multiome library construction and data analysis

#### step1, cell preparation

For v2a-v2b lineages, single-cell suspensions were prepared and GFP-high cells of *Tg(vsx1:GFP)* were isolated by fluorescence-activated cell sorting (FACS) as described for the scRNA-seq analysis above. For AC-BC lineages, cells were prepared as previous published1(*3*). Briefly, single-cell suspensions were prepared from dissected retinas of *Tg(atoh7:gapRFP::atoh7:turboGFP-dest1)* embryos at 44 hpf, GFP^+^RFP^-^ cells were subsequently collected by FACS.

#### Step2, nuclei isolation

The FACS-sorted cell suspension was centrifuged at 500 g for 5 minutes 4 °C. The supernatant was discarded. The cell pellet was resuspended in 50 μl PBS-0.04% BSA and centrifuged at 300 g for 5 minutes 4 °C. After removing 45 μl of supernatant, 45 μl of 0.5x chilled lysis buffer (10 mM Tris-HCl, 10 mM NaCl, 3 mM MgCl2, 0.05% Tween-20, 0.05% NP40, 0.005% Digitonin, and 1% BSA, 1 mM DTT, 1 U/μl RNase inhibitor) was added. The sample was gently pipetted three times and incubated on ice for 3 minutes.

Subsequently, 50 μl of chilled wash buffer (10 mM Tris-HCl, 10 mM NaCl, 3 mM MgCl2, 0.1% Tween-20, and 1% BSA, 1 mM DTT, 1 U/μl RNase inhibitor) was added without pipetting. The sample was immediately centrifuged at 500 g for 5 minutes at 4 °C. After removing 95 μl of supernatant, 45 μl of ice-cold Diluted Nuclei Buffer (10x Genomics) was added without pipetting and centrifuged again at 500 g for 5 minutes at 4 °C. Finally, 45 μl of supernatant was removed and the nuclei pellet was resuspended in 5 μl of chilled diluted nuclei buffer.

#### Step3, library construction and sequencing

Approximately 10,000 nuclei were loaded for simultaneous single-nucleus RNA sequencing (snRNA-seq) and single-cell ATAC sequencing (scATAC-seq) using the Chromium Single Cell Multiome ATAC + Gene Expression kit (10x Genomics, Cat# PN-1000283) according to the manufacturer’s protocol. The snRNA-seq (Gene Expression, GE) library was sequenced on an Illumina NovaSeq 6000 platform with 150 bp paired-end (PE150) reads. The scATAC-seq (ATAC) library was sequenced on an MGISEQ-2000 platform with 100 bp paired-end (PE100) reads.

#### Step4, data processing

The ATAC sequencing Read1 and Read2 FASTQ files were split into four files (R1, R2, R3, I1). Both GE and ATAC FASTQ files were renamed, aligned to the zebrafish reference genome (GRCz11), filtered, and quantified using Cell Ranger ARC software (v2.0.1, 10x Genomics) following the manufacturer’s instructions. The downstream analysis was carried out using the R package Seurat (v5.1.0) and Signac (v1.13)(*42*). Data integration and multimodal clustering were achieved using Weighted Nearest Neighbor (WNN) analysis (*43*).

### In silicon analysis of *tal1* and ptf1α upstream repressor candidates

Cis-regulatory elements within the tal1 and ptf1α promoter were manually annotated. Genome sequences of these CREs were retrieved from Ensembl (https://www.ensembl.org/). Potential upstream regulators binding to these CREs were predicted using the PredictTFBS function in AnimalTFDB4 (https://guolab.wchscu.cn/AnimalTFDB4)(44). Transcriptional profiles of predicted transcription factors (TFs) across cell states were extracted from our scRNA-seq data. TFs with average expression > 0.15 (log-normalized counts) in precursor cells were selected for downstream analysis. Candidate repressors were identified by comparing TF expression in excitatory (v2a or BC) versus inhibitory (v2b or AC) neuronal lineages, retaining factors meeting both criteria: 1, Significantly higher expression in excitatory neurons (p < 0.05); 2, Average fold-change > 2. Homeobox TF motif logos were obtained from the CIS-BP Database (https://cisbp.ccbr.utoronto.ca)

### CUT&Tag library construction and data analysis

Since her15.1 is widely expressed in whole body, we cross *Tg(lhx3:gal4)* with *Tg(uas:her15.1-v5-p2a-tdTomatoNLS)* to restrict her15.1 profiling in ventral spinal cord. At 24 hpf, embryos exhibiting tdTomato fluorescence were selected. Heads and yolk sacs were removed, and 50 trunks were dissociated into single-cell suspensions at room temperature. For hey1 study, *TgKI(hey1-v5-p2a-gal4)* was crossed with *Tg(uas:kaede)*. About 30 retinas were dissected from 48 hpf F1 embryos and dissociated into single-cell suspensions at room temperature. Freshly prepared single-cell suspensions were immediately processed using commercial CUT&Tag Assary Kit (Vazyme, Cat# TD-903) according to the manufacture’s protocol. Briefly, cells were immobilized by Concanavalin A-coated beads, sequentially incubated with: mouse-anti-v5 antibody (1:50, Bio-rad, Cat# MCA1360GA) overnight at 4 °C, rabbit-anti-mouse IgG (1:100, Abcam, Cat# Ab46540) and Protein A/G-pTn5 transposase complex; followed by fragmentation, DNA extraction and PCR amplification (20 cycles for *Her15.1*, 17 cycles for *Hey1*). Libraries were sequenced on Illumina NovaSeq 6000 (2×150 bp). Sequenced results were processed using CUT&RUNTools2.0 pipline (*45*), adapted for zebrafish (GRCz11). Analysis includes read trimming, alignment (GRCz11), BAM processing, peak calling (MACS2 and SEACR), motif finding and motif footprinting (macs2.narrow.120). Peaks presented in this work were called from MACS2 (*46*).

### LY411575 treatment to block Notch signaling pathway

Apply 10 μM LY411575 (Selleck Chemical, Cat# S2714) or 10 μM DMSO to *TgKI(hey1^mneogreen/+^)* in the embryo medium at 24 hpf, take image at 48 hpf.

### Image processing

Image analysis was performed with FV10-ASW 4.0 (Olympus), and Image J. For presentation, maximal intensity projection of several z-sections was used, brightness and contrast of each channel were adjusted separately. All manipulations were applied for the whole pictures, and then the regions of interest were selected and cropped from the whole picture for presentation.

### Quantification and statistical analysis

All cell types and cell numbers were determined manually. When studying effects of TF overexpression, we compare the difference between two groups (e.g., blank vs tal1 OE) with the sum of all collected cells, rather than the quantification fish by fish. We decided this kind of quantification because the cells with TF overexpression distribute sparsely and variably in spinal cords. χ² test was used for significant difference determination in this situation. Cell numbers of each group are provided in the legends. Cells were collected from more than 10 fish for v2a-v2b lineage study, more than 7 fish for AC-BC lineage study. The input data to do χ² test is like the following table:

|  | blank | tal1 OE |
| --- | --- | --- |
| gad1b+ | 77 | 111 |
| gad1b- | 172 | 84 |

For t-test in other situations, the normality assumption of the data was tested with first. For two-group comparison, the two-tailed unpaired *t*-test was used for significance analysis for normal data. Sample sizes, error bars, and exact *p* values are provided in figures and legends. Analysis was performed using GraphPad Prism v8.0 software. Data are mean± SD.

---

**Fig. S1.**
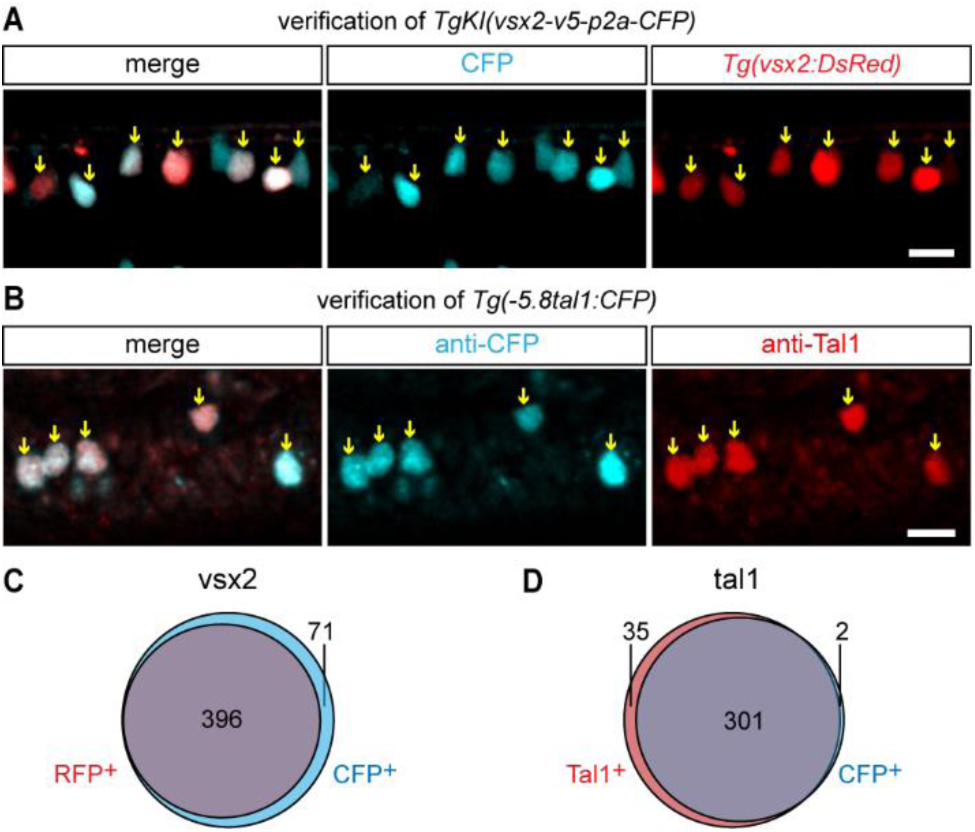
Validation of v2a and v2b reporter fishlines. (**A**) Cross of *TgKI(vsx2-v5-p2a-CFP)* and established vsx2 reporter *Tg(vsx2:Dsred)*. (**B**) Double immunostaining of CFP and Tal1 in *Tg(−5.8tal1:CFP)*. (**C**) Quantification of reporter overlap from (A). (**D**) Quantification of Tal1 and CFP colocalization from (B). Cell numbers indicated. Scale bars: 10 μm.

**Fig. S2.**
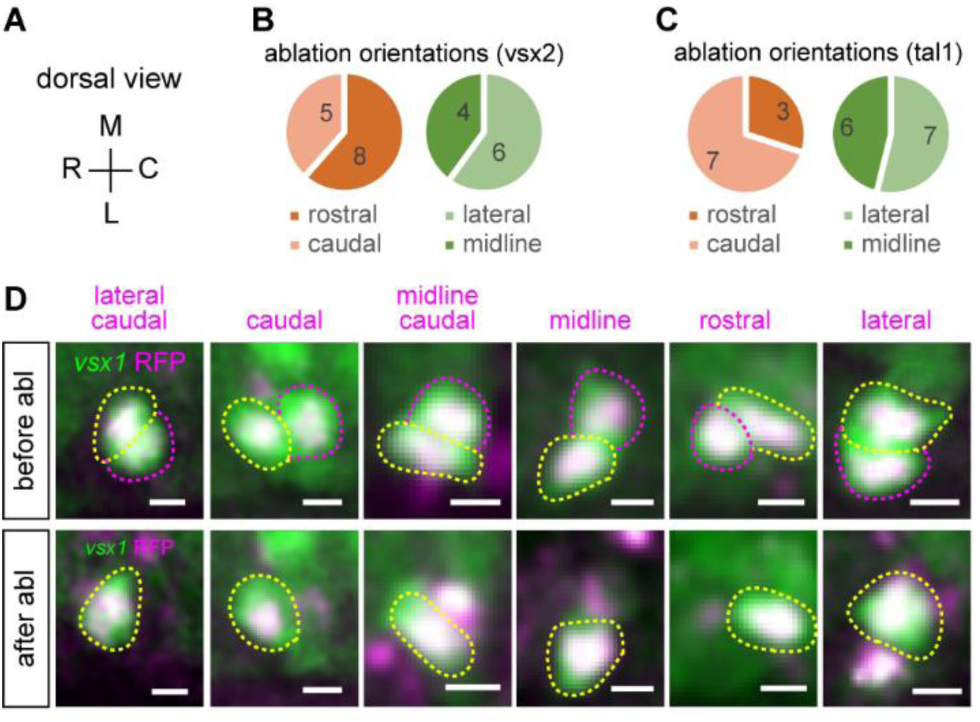
Orientations of sibling cell ablations. (**A**) Schematic of fish orientation from dorsal view. (**B** and **C**) Ablation distribution along rostral-caudal and lateral-midline axis in *vsx2* (B) and *tal1* (C) reporter lines. (**D**) Representative ablations in different orientations. Cell indicated by magenta circle is the ablation target. Scale bars: 5 μm.

**Fig. S3.**
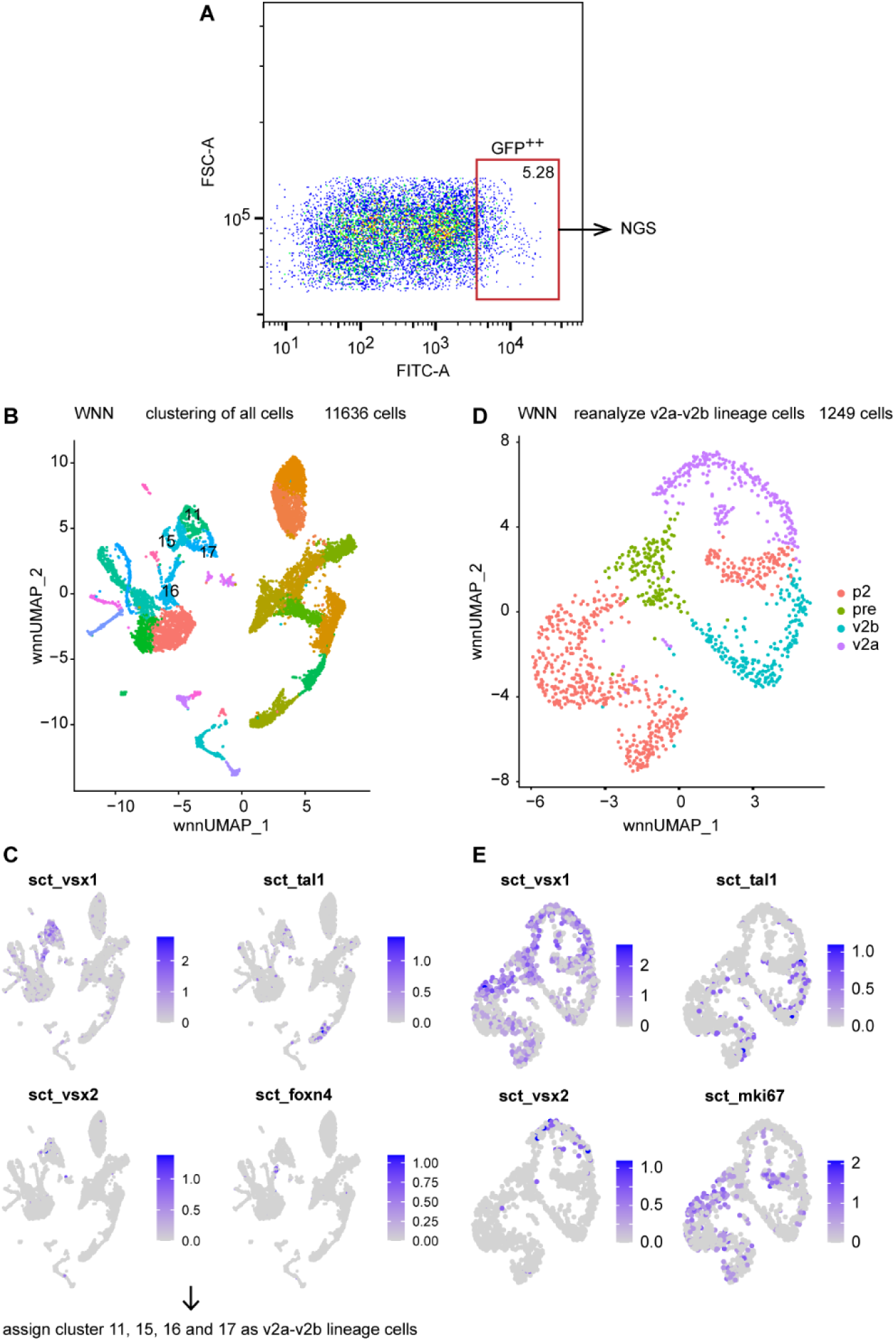
Single-cell multiome analysis of v2a-v2b lineages. (**A**) Sample preparation: FACS sorting of GFP^++^ cells from *Tg(vsx1:GFP)*. (**B**) Integrated RNA + ATAC clustering of all sequenced cells. (**C**) V2a-v2b lineage marker expression across clusters. Cells of cluster 11, 15, 16 and 17 in (B) are assigned as v2a-v2b lineage cells. (**D**) Re-clustering of v2a-v2b lineage cells. (**E**) Expression of stage-specific markers. P2: *mki67*; v2a: *vsx2*; v2b: *tal1*; pre: intermediate state.

**Fig. S4.**
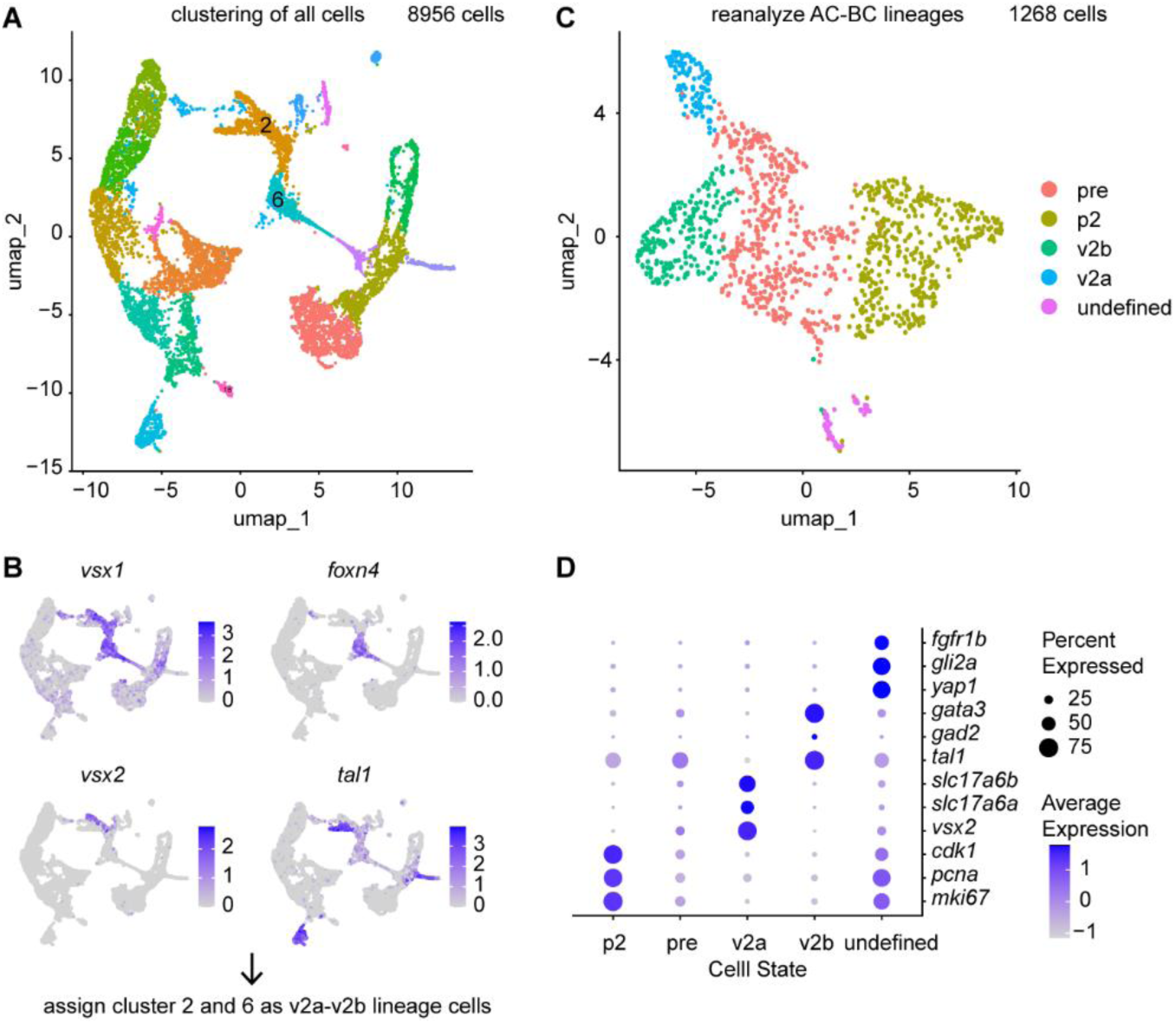
Single-cell RNA sequencing analysis of v2a-v2b lineages. (**A**) Clustering of all sequenced cells. (**B**) V2a-v2b lineage marker expression across clusters. Cells of cluster 2 and 6 in (A) are assigned as v2a-v2b lineage cells. (**C**) Re-clustering of v2a-v2b lineage cells. (**D**) Expression of stage-specific markers.

**Fig. S5.**
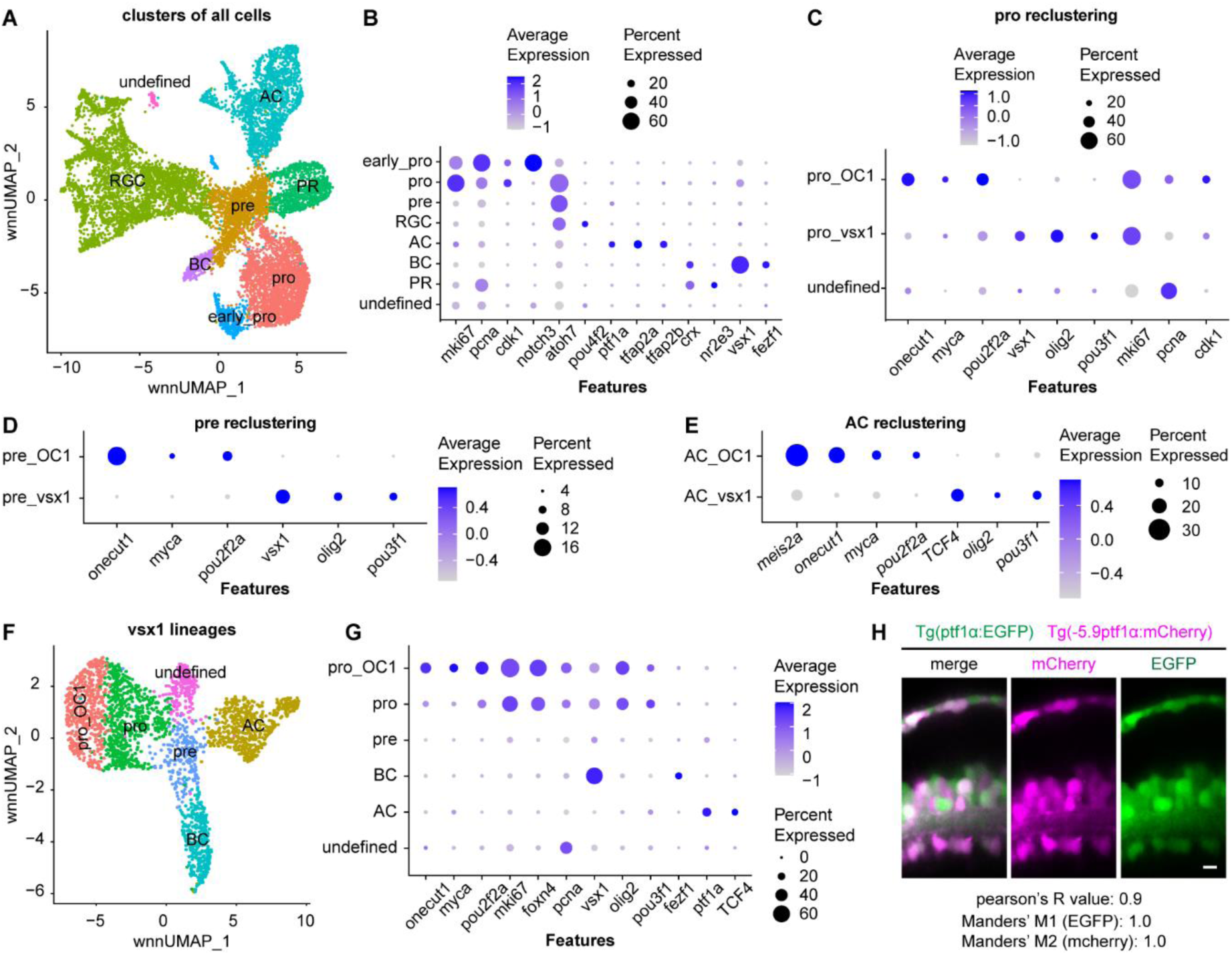
Single-cell multiome analysis of of AC-BC lineages. (**A** and **B**) Clustering of all sequenced cells with RNA + ATAC integration: (A) Cell clusters; (B) cell marker expression. (**C**-**E**) Re-clustering of: (C) progenitors; (D) Precursors; (E) ACs. (**F** and **G**) Re-analysis of all *vsx1* lineage cells. *Vsx1* lineage cells are the combining of annotated BC, pro_*vsx1*, pre_*vsx1*, AC_*vsx1* clusters. (F) New clustering; (G) Cell marker expression. (**H**) Correlation between the reporter signals driven by 5.9 kb upstream of *ptf1α* and *ptf1α* BAC promoter.

**Fig. S6.**
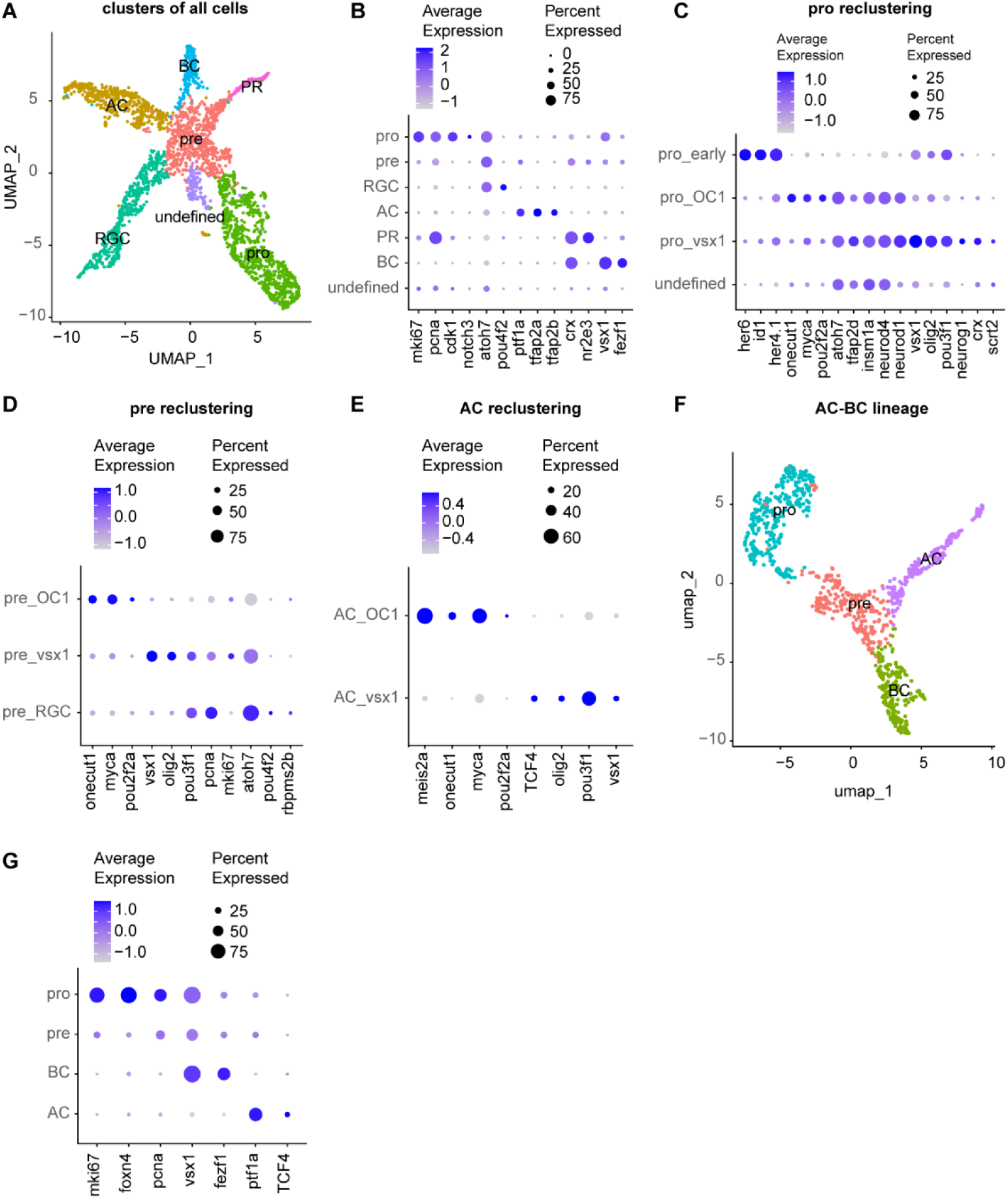
ScRNAseq analysis of AC-BC lineages. (**A** and **B**) Clustering of all sequenced cells: (A) Cell clusters; (B) cell marker expression. (**C**-**E**) Re-clustering of: (C) Progenitors; (D) Precursors; (E) ACs. (**F** and **G**) Re-analysis of *vsx1* lineage cells. *Vsx1* lineage cells are the combining of annotated BC, pro_*vsx1*, pre_*vsx1*, AC_*vsx1* clusters. (F) New clustering; (G) Cell marker expression.

**Table S1.**
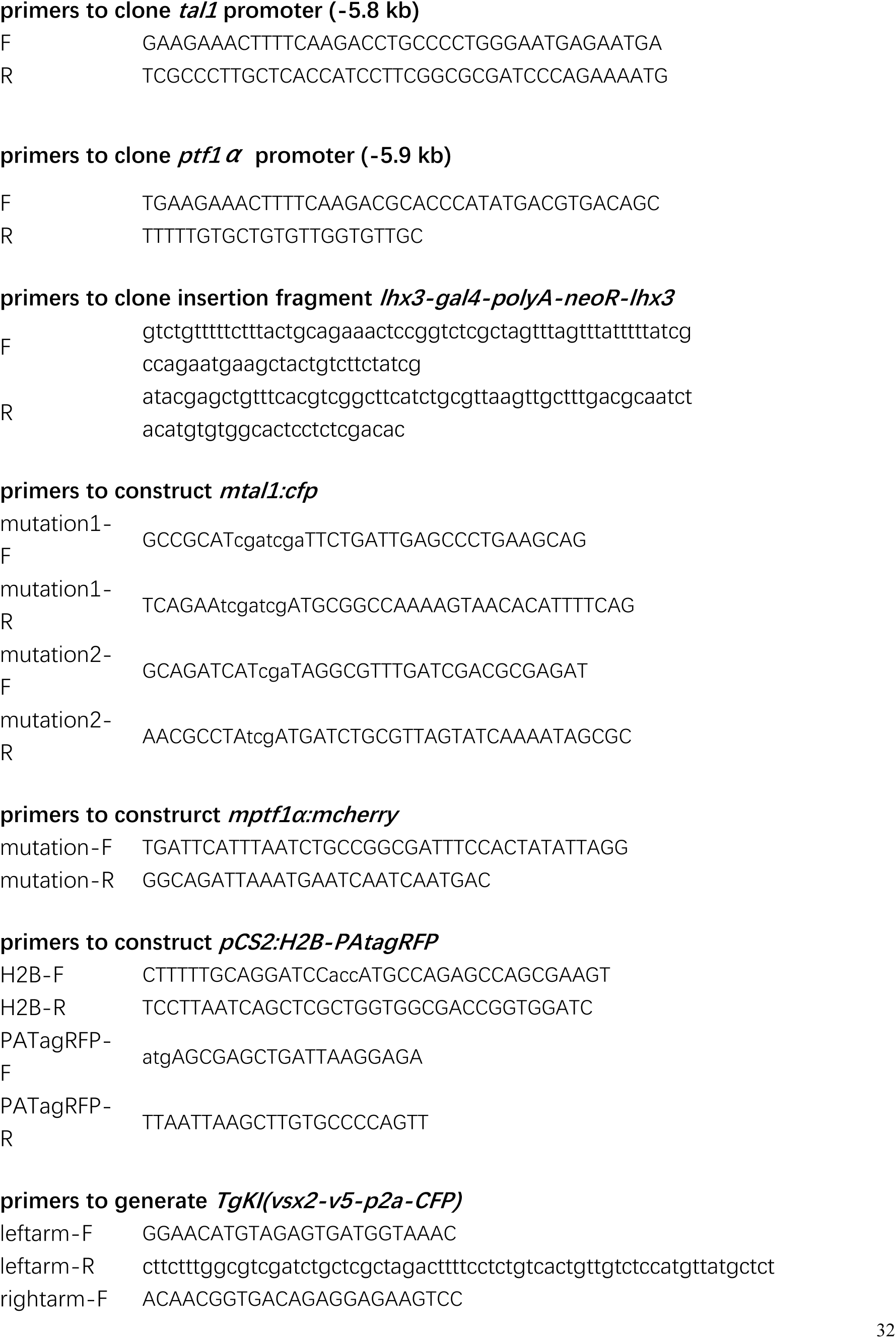
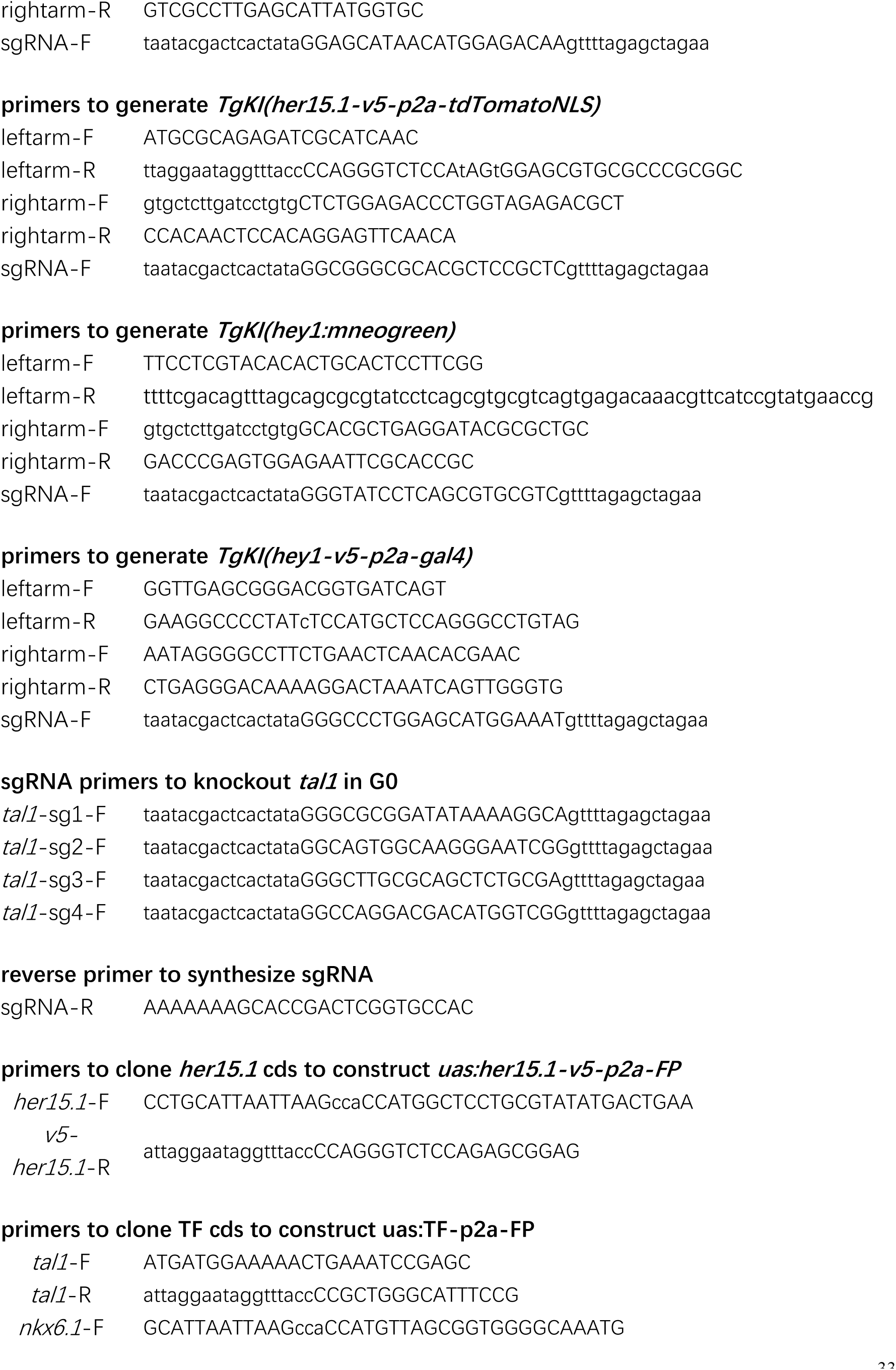
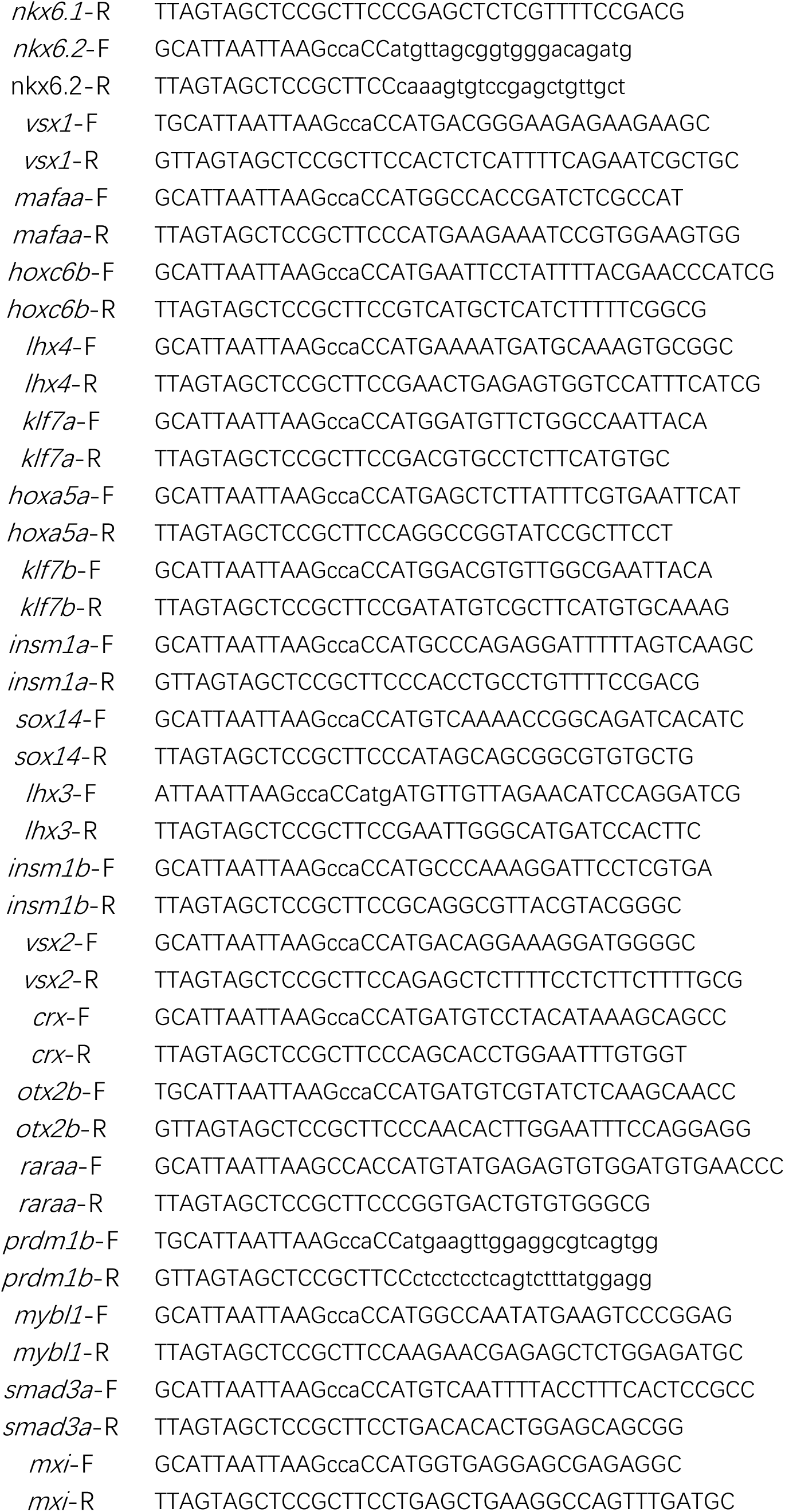
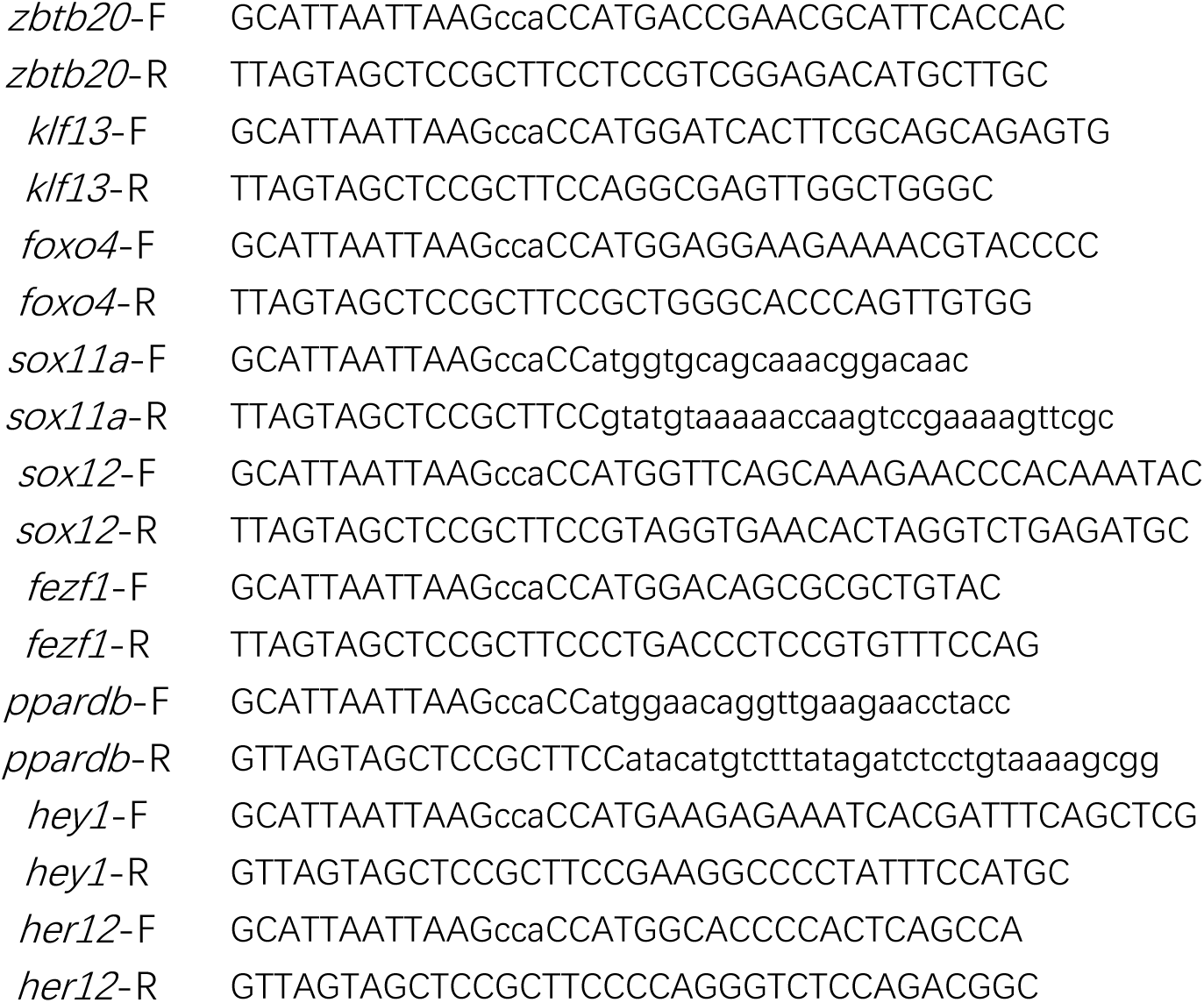
Primers.

